# Like sisters but not twins – vasopressin and oxytocin excite BNST neurons via cell type-specific expression of oxytocin receptor to reduce anxious arousal

**DOI:** 10.1101/2024.09.06.611656

**Authors:** Walter Francesconi, Valentina Olivera-Pasilio, Fulvia Berton, Susan L. Olson, Rachel Chudoba, Lorena M. Monroy, Quirin Krabichler, Valery Grinevich, Joanna Dabrowska

## Abstract

Interoceptive signals dynamically interact with the environment to shape appropriate defensive behaviors. Hypothalamic hormones arginine-vasopressin (AVP) and oxytocin (OT) regulate physiological states, including water and electrolyte balance, circadian rhythmicity, and defensive behaviors. Both AVP and OT neurons project to dorsolateral bed nucleus of stria terminalis (BNST_DL_), which expresses oxytocin receptors (OTR) and vasopressin receptors and mediates fear responses. However, understanding the integrated role of neurohypophysial hormones is complicated by the cross-reactivity of AVP and OT and their mutual receptor promiscuity. Here, we provide evidence that the effects of neurohypophysial hormones on BNST excitability are driven by input specificity and cell type-specific receptor selectivity. We show that OTR-expressing BNST_DL_ neurons, excited by hypothalamic OT and AVP inputs via OTR, play a major role in regulating BNST_DL_ excitability, overcoming threat avoidance, and reducing threat-elicited anxious arousal. Therefore, OTR-BNST_DL_ neurons are perfectly suited to drive the dynamic interactions balancing external threat risk and physiological needs.

**Graphical abstract:** 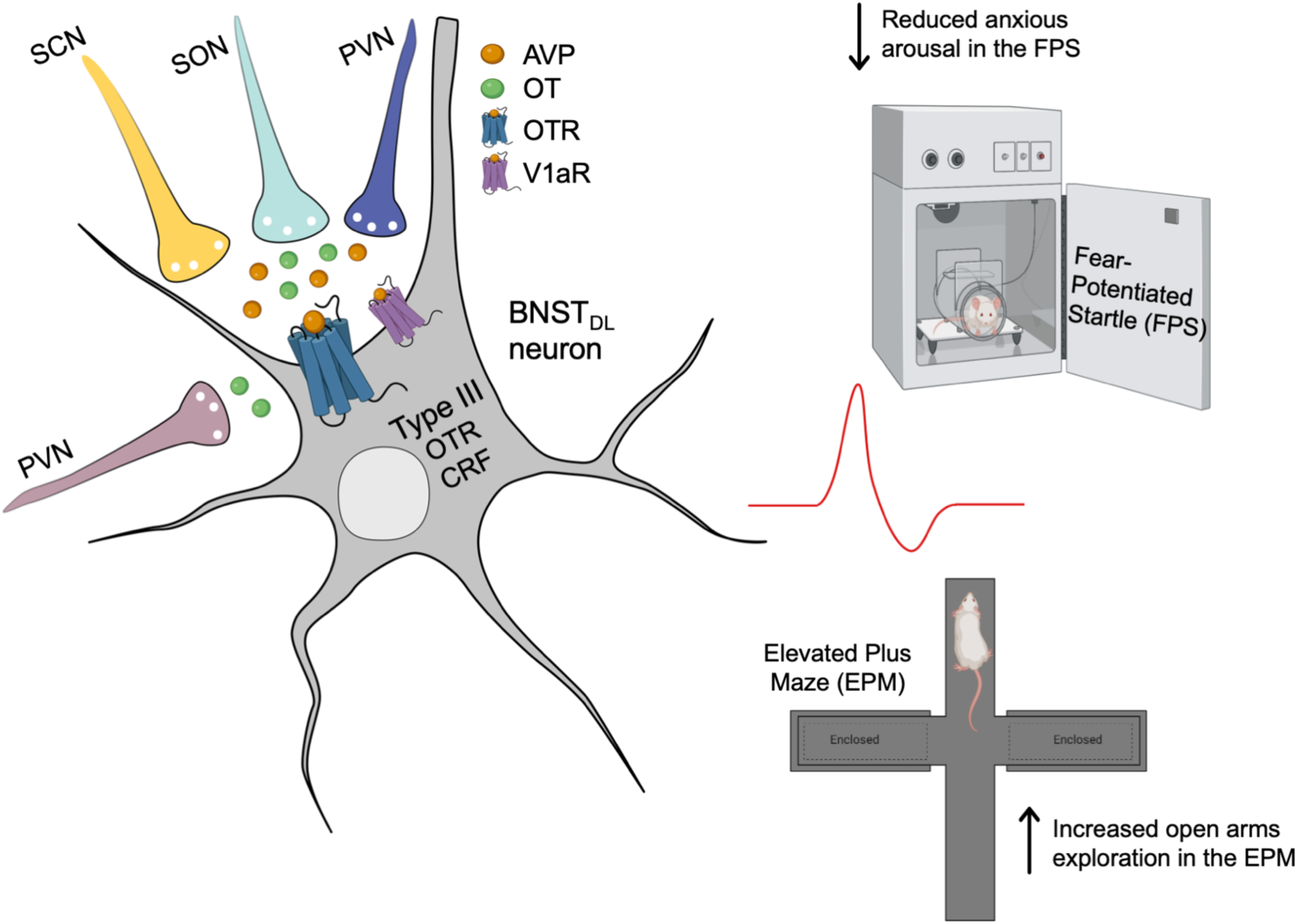

**Highlights:**

- Exogenous and light-evoked vasopressin (AVP) peptide excites neurons of the bed nucleus of the stria terminalis (BNST) via oxytocin receptor (OTR) in male rats
- AVP excites OTR- and Corticotropin-releasing factor (CRF)-expressing neurons, most of which are classified as Type III neurons of the BNST
- OTR-expressing BNST neurons increase exploration of open spaces and reduce anxious arousal in fear-potentiated startle in male rats
- The dorsolateral BNST receives vasopressinergic inputs from suprachiasmatic, supraoptic, and paraventricular nuclei of the hypothalamus
- Internal signal-sensitive hypothalamic inputs directly impact BNST excitability via OTR to balance interoceptive signals and defensive behaviors

## 1. Introduction

Interoceptive signals dynamically interact with the environment to shape appropriate defensive behaviors. For example, rodents avoid open spaces to escape from predators but must overcome this defensive behavior to locate water when driven by thirst. Further, circadian rhythm-associated interoceptive signals drive animals to follow wake-sleep cycles despite predator threats. Numerous physiological functions, including water and electrolyte balance and circadian rhythmicity, are supported by the hormone and neuromodulator arginine-vasopressin (AVP) ^1,2^, making AVP a prime candidate for modulating the interaction balancing external risks and internal needs. However, it is not clear how AVP acts in the extended amygdala, which directly influences defensive responses, to shape appropriate, physiologically relevant levels of exploratory and defensive behaviors.

AVP signals via three centrally located receptors: vasopressin receptors (V1aR, V1bR) and the oxytocin receptor (OTR), all of which are implicated in defensive behaviors, including fear, avoidance- and anxiety-like behaviors, for review see ^3–8^ (of note V2R is located in the kidneys ^9^). The receptors are differentially expressed in the brain, with V1bR highly expressed in the pituitary and other discrete locations ^10,11^, whereas V1aR and OTR more highly expressed throughout the central nervous system ^12–14^. Notably, OTR and V1aR seemingly have opposite effects on exploration and fear-related behaviors in rodents ^15,16^, suggesting that receptor selectivity is one mechanism by which AVP modulates defensive behaviors. The extended amygdala, including the central (CeA) and medial amygdala (MeA), as well as bed nucleus of the stria terminalis (BNST), has a high density of both oxytocin (OT)- and AVP-binding sites ^17^. However, specific receptor contributions to behavioral and physiological responses remain unclear and likely underlie the diverse behavioral effects of AVP. For example, there is a mismatch between AVP peptide density and vasopressin receptor expression in some regions of the extended amygdala, such as sparse V1aR but high AVP fiber density and OTR binding sites in posterior BNST, for review see ^18^. In addition, OTR contribution to AVP signaling may be obscured in behavioral studies using prominent V1R antagonists that also bind OTRs ^19,20^. Given the abundance of OTRs and V1Rs in hypothalamic and the extended amygdala, AVP’s effects via OTR are likely to be vastly underestimated.

Besides receptor selectivity, the diverse effects of AVP on behavior may also be modulated by input specificity. AVP is produced in neurons of the hypothalamus–including the paraventricular (PVN), supraoptic (SON), and suprachiasmatic nucleus (SCN) ^21,22^–and the extended amygdala– including the MeA and posterior BNST ^23^. As AVP and OT predominantly exist in separate neuronal groups within the hypothalamus ^24^, AVP and/or OT inputs, prompted by specific physiological cues, might orchestrate the balance between physiological states and defensive behaviors through downstream brain structures. Given the hypothalamus’ role in assessing internal physiological states and the extended amygdala’s role in gauging external threats, AVP and OT likely flexibly regulate defensive behaviors based on the interplay between physiological needs and external threats.

The dorsolateral portion of the BNST (BNST_DL_) is critical for fear processing and vigilant threat monitoring ^25–28^ and is one of the few brain regions with high expression of OT and AVP binding sites and fibers ^17,29^, as well as OTR and V1aR ^30–32^. It receives inputs from hypothalamic OT (primarily PVN) and AVP neurons ^23,33,34^, but the origins of the AVP projections to the BNST_DL_ are unknown. Primarily a GABA-ergic and peptidergic structure ^35,36^, the BNST_DL_ contains three major neuron types (Type I-III) defined by their intrinsic electrophysiological membrane properties and firing patterns ^37,38^. Previously, we demonstrated that OT is released in the BNST_DL_ in response to fear stimuli ^39^ where it has distinct cell type-specific effects on neuronal excitability and synaptic transmission ^40^. However, the integrative mechanisms by which AVP and OT act via V1R and/or OTR to regulate neuronal excitability and fear processing in the BNST_DL_ are unknown.

In this study, we used slice electrophysiology, neuronal tract tracing, and peptide optogenetics in naïve and AVP-Cre transgenic male rats to show that AVP neurons from the SCN, SON, and PVN project to the BNST_DL_. Both exogenous and blue light-evoked AVP excited Type I and Type III BNST_DL_ neurons, which required OTR transmission. In contrast, V1aR produced varying excitatory or inhibitory effects depending on neuron type. Using OTR-Cre transgenic rats ^41,42^, we confirmed that fluorescent OTR-BNST_DL_ neurons, which match the properties of Type I and Type III cells, were excited by AVP. Single-cell PCR previously showed that Type III BNST_DL_ neurons express both OTR and CRF mRNA ^33^. Using CRF-Cre transgenic rats ^43^, we demonstrated that AVP and an OTR agonist (TGOT) directly excited these CRF-BNST_DL_ neurons. Finally, using chemogenetics, we show that OTR-BNST_DL_ neurons’ activity reduced anxious arousal in the fear-potentiated startle and increased exploration in the elevated-plus maze. Our findings show that AVP and OT effects on BNST_DL_ excitability depend on cell-type specific receptor expression and hypothalamic input specificity. Therefore, changes in the activity of internal signal-sensitive hypothalamic nuclei will directly impact BNST_DL_ excitability to balance exploratory and defensive behaviors.

## 2. Star Methods

### Experimental models and subjects

Male wild-type Sprague-Dawley rats (SD, Envigo, Chicago, IL; 240-300 g), male transgenic OTR-Cre (Cre-recombinase (Cre) under OTR promoter), CRF-Cre (Cre under CRF promoter), and AVP-Cre (Cre under AVP promoter) rats (220-400 g) were housed in groups of three on a 12-h light/dark cycle (light 7 a.m. to 7 p.m.) with free access to water and food. Upon arrival from Envigo, wild-type SD rats were habituated to the environment for one week before the experiments began. OTR-Cre ^41,42^ and AVP-Cre transgenic (knock-in) rats were originally generated and kindly provided by Dr. Valery Grinevich (Heidelberg University, Germany) ^44^, and CRF-Cre rats ^45,46^ were created and kindly provided by Dr. Robert Messing (University of Texas, Austin) and bred at the biological resources facility (BRF) at Rosalind Franklin University of Medicine and Science (RFUMS). Experiments were performed in accordance with the guidelines of the NIH and approved by the Animal Care and Use Committee at RFUMS.

### Drug preparation and pharmacological manipulations

The following compounds were used for electrophysiology: arginine (Arg^8^)-vasopressin (AVP; H-Cys-Tyr-Phe-Gln-Asn-Cys-Pro-Arg-Gly-NH₂ (disulfide bond); 0.2 μM, Bachem Americas, Torrance, CA; catalog #4012215); oxytocin (OT; H-Cys-Tyr-Ile-Gln-Asn-Cys-Pro-Leu-Gly-NH2 (disulfide bond, acetate salt), 0.2 μM, Bachem Americas, Torrance, CA, #4016373); a selective rat V1aR agonist FE201874 (0.2-0.4 μM, generously provided by Ferring Pharmaceuticals, San Diego, CA) ^47^, a selective and potent OTR agonist TGOT ([Thr^4^,Gly^7^]-oxytocin (0.4 μM, Bachem Americas, Torrance, #4013837), and V1bR agonist d[Cha4]-AVP (1 μM, GlpBio Technology Inc, Montclair, CA, #GC1659) ^3,48^. The following antagonists were used: a V1aR/OTR antagonist (d(CH_2_)_5_^1^,Tyr(Me)^2^,Arg^8^)- vasopressin, Manning compound ^49^; 1 μM, Tocris, Bio-Techne Corporation; MN, #3377); selective OTR antagonist (OTA; d(CH2)5(1), D-Tyr(2), Thr(4), Orn(8), des-Gly-NH2(9)]-Vasotocin trifluoroacetate salt ^49^; 0.4 μM, Chemical Repository, #V-905 NIMH); selective V1aR antagonist SR49059 (2*S*)-1-[[(2*R*,3*S*)-5-Chloro-3-(2-chlorophenyl)-1-[(3,4-dimethoxyphenyl)sulfonyl]-2,3-dihydro-3-hydroxy-1*H*-indol-2-yl]carbonyl]-2-yrrolidinecarboxamide; 5 μM, Tocris, Bio-Techne Corporation, #2310); and a V1bR antagonist Nelivaptan ((2*S*,4*R*)-1-[(3*R*)-5-Chloro-1-[(2,4-dimethoxyphenyl)sulfonyl]-2,3-dihydro-3-(2-methoxyphenyl)-2-oxo-1*H*-indol-3-yl]-4-hydroxy-*N*,*N*-dimethyl-2-pyrrolidinecarboxamide; 1 μM, Tocris, Bio-Techne Corporation, #6195). To block glutamatergic transmission, we used AMPA and kainate receptor antagonist 6-Cyano-7-nitroquinoxaline-2,3-dione disodium salt (CNQX, 10 μM, Tocris, Bio-Techne Corporation, #1045). To block NMDA receptors, we used D-2-Amino-5-phosphopentanoic acid (D-AP5, 50 μM, Tocris, Bio-Techne Corporation, #0106). To block GABA-A receptors we used picrotoxin (PTX, 25 μM, Tocris, Bio-Techne Corporation, #1128). The designer receptor exclusively activated by designer drugs (DREADD) ligand clozapine-N-oxide (CNO), 8-Chloro-11-(4-methyl-4-oxido-1-piperazinyl)-5*H*-dibenzo[*b*,*e*][1,4]diazepine (CNO, Tocris, Bio-Techne Corporation, #4936) was used at 20 μM for chemogenetics manipulations during electrophysiological recordings and at 2 mg/kg for behavioral experiments. All drug stock solutions were dissolved in sterile deionized water, unless stated otherwise, and stored at −20°C until use. The day of the experiment, drugs for electrophysiology were diluted into artificial cerebrospinal fluid (aCSF) and applied to bath solution. CNO was diluted into sterile saline and injected intraperitoneally (i.p.) at 2 mg/kg.

### *In vitro* whole-cell patch-clamp electrophysiology

Slice preparation and electrophysiological recordings were performed as before ^40^. Rats were deeply anesthetized by inhalation of isoflurane USP (Patterson Veterinary, Greeley, CO, USA). After decapitation, the brain was rapidly removed from the cranial cavity, and 300-μm-thick coronal slices containing the BNST were prepared in ice-cold cutting solution (saturated with 95% O_2_/ 5% CO_2_) containing in mM: 122.5 NaCl, 3.5 KCl, 25 NaHCO_3_, 1 NaH_2_PO_4_, 0.5 CaCl_2_·2H_2_O, 20 D-glucose, 3 MgCl_2_ 6H_2_O, and 1 ascorbic acid (pH 7.4, 290–300 mOsm). The slices were prepared using a Leica vibratome (VT1200; Leica, Wetzlar, Germany), incubated for 30 min at 34°C, and subsequently transferred to room temperature for 1 hour before the recordings began. Next, the slices were transferred to a recording chamber perfused with oxygenated aCSF at a rate of 2-4 ml/min containing in mM: 122.5 NaCl, 3.5 KCl, 25 NaHCO_3_, 1 NaH_2_PO_4_, 2.5 CaCl_2_·2H_2_O, 20 D-glucose, and 1 MgCl_2_·6H_2_O (pH 7.4, 290-300 mOsm), saturated with 95% O_2_ and 5% CO_2._ The aCSF was warmed to 30–34°C by passing it through a feedback-controlled in-line heater (TC-324C; Warner Instruments, Hamden, CT). The cell bodies of BNST_DL_ neurons were visualized using infrared differential interference contrast (IR-DIC) optics with an upright microscope (Scientifica Slice Scope Pro 1000, Clarksburg, NJ). Neurobiotin (0.1%, Vector Laboratories, Burlingame, CA) was added to the internal solution of the recording pipette to confirm cell location. After recording, slices were immersed in 10% formalin (Fisher Scientific, SF98-4), washed three times (10 min each) in phosphate-buffered saline (PBS 0.05 M), and incubated with streptavidin-Alexa Fluor 594 or Alexa Fluor 488 conjugate (dilution 1:2000, Invitrogen ThermoFisher Scientific, S32356 and S11223, respectively) at room temperature for 2 hours, followed by three washes in PBS and one wash in phosphate buffer (PB 0.05 M) (10 min each). All slices were mounted using Mowiol with antifade reagent (Sigma-Aldrich, 81381), and coverslips were applied before visualization with fluorescent microscopy (Nikon eclipse N*i*, Nikon Instruments Inc.)

Electrophysiology recordings were made using Multiclamp 700B amplifiers (Axon Instruments, Union City, CA); current-clamp signals were acquired at 10 kHz with a 16-bit input-output board NI USB-6251 (National Instruments, Austin, TX) using custom MatLab scripts (Math-Work, Natick, MA) written by Dr. Niraj Desai ^50^. The access resistance (Ra) was monitored, and recordings were terminated if Ra changed > 15%. Electrophysiological measurements were carried out 10-15 min after reaching the whole-cell configuration. The input resistance was calculated from steady-state voltage responses upon negative current injections (pulses of 450-1000 ms). Whole-cell patch-clamp recordings were made from BNST_DL_ neurons using glass pipettes (4–8 MΩ) pulled from thick-walled borosilicate glass capillaries with a micropipette puller (Model P-97; Sutter Instrument, Novato, CA) filled with a solution containing the following in mM: 135 potassium gluconate, 2 KCl, 3 MgCl2.6H_2_O, 10 HEPES, 5 Na-phosphocreatine, 2 ATP-K, and 0.2 GTP-Na (pH 7.3 adjusted with KOH, osmolarity 300-305 mOsm; potentials were not corrected for a liquid junction potential), as before ^40^.

To study the effects of AVP on intrinsic membrane properties and intrinsic excitability, the recordings were performed in the presence of synaptic transmission blockers: the AMPA receptor antagonist CNQX, the NMDA receptor antagonist D-AP5, and GABA-A receptor antagonist, PTX. To verify the receptor involved in the effects of AVP, in separate experiments, AVP was perfused in the presence of the V1aR/OTR antagonist, selective OTR antagonist OTA, selective V1aR antagonist SR49059, or selective V1bR antagonist Nelivaptan. Each antagonist was first applied alone for 10 min and then maintained during AVP application for another 12-15 min, followed by a 12-15-min drug-free washout in all whole-cell patch-clamp experiments.

### Electrophysiological characterization of BNST_DL_ neurons

At the beginning of the recording sessions in current clamp mode, neurons were characterized using current pulses (450 ms in duration) from −250pA to 180pA in 10-pA increments. Based on their characteristic voltage responses ^37^, three major types of neurons were identified in the BNST_DL_. Type I neurons fire regularly and display moderate spike frequency adaptation. These neurons also display a voltage sag that indicates the presence of the hyperpolarization-activated cation current (I_h_). As a distinguishing feature of Type II neurons, post-inhibitory spikes are produced in response to preceding negative current steps that are related to the action of the low-threshold Ca^2+^ current. Additionally, Type II neurons display strong voltage sags under hyperpolarizing current pulses that indicate a high level of I_h_ current. Type III neurons differ from the previous two types in several aspects: they exhibit high rheobase and no voltage sag under negative current steps; they display prominent inward rectification that is caused by the activation of the inward rectifying K^+^ current (I_KIR_) at membrane potentials more negative than approximately −50 mV; and start firing after a characteristic slow voltage ramp mediated by the K^+^ delayed current (I_D_).

### The effect of AVP on membrane properties and intrinsic excitability of Type I, II, and III BNST_DL_ neurons

We investigated the effects of AVP and other pharmacological manipulations in Type I-III BNST_DL_ neurons on the following membrane properties: resting membrane potential (RMP), input resistance (Rin), rheobase (Rh), threshold of the first action potential (first-spike Th) (calculated as the voltage at which the depolarization rate exceeded 5 mV/ms), and latency of the first spike (first-spike Lat) evoked at the Rh current. We also investigated the intrinsic excitability by measuring the input-output (I/O) relationship before (pre), during, and after AVP application (post). We measured the steady-state frequency (SSF) by applying depolarizing pulses (1 sec in duration) of different amplitudes (0-180 pA in 10 pA increments) and determining the average of the inverse of the inter-spike intervals (ISI) from all action potentials starting from the second action potential. We used SSF as a function of current to assess the effect of AVP on intrinsic excitability of Type I-III BNST_DL_ neurons from wild-type rats, as well as fluorescent BNST_DL_-OTR neurons and BNST_DL_-CRF neurons, from OTR-Cre and CRF-Cre transgenic rats, respectively.

### Electrophysiological recordings from fluorescent neurons

To visualize OTR- and CRF-mCherry fluorescent neurons in brain slices from OTR-Cre and CRF-Cre rats or AVP-eYFP neurons in the hypothalamus (SON, SCN, and the PVN), we used an upright microscope (Scientifica Slice Scope Pro 1000 fitted with fluorescent filters (49008_Olympus BX2_Mounted, ET mCherry, Texas Red, ET560/40x ET630/75m T585lpxr for the visualization of mCherry and 49002 ET-EFP (FITC/Cy2) ET470/40x ET525/50m T495LPXR for the visualization of eYFP) and infrared differential interference contrast [IR-DIC] optics), with a CoolLED pE-300^ultra^, broad-spectrum LED illumination system as the light source. We identified fluorescent neurons using the 40x objective, mCherry or eYFP filter, and a live-image video camera (IR-2000, Dage-MTI). Once these neurons were targeted and considered healthy using an IR-DIC filter, we performed the whole-cell patch-clamp recordings.

### Stereotaxic surgeries and adeno-associated virus (AAV) injections

We deeply anesthetized male OTR-Cre rats (n=92), Wistar CRF-Cre rats (n=26), both at approximately 250g of body weight, +/-10 g), and AVP-Cre rats (n=83, at 220g (+/-10 g) body weight) with a mix of isoflurane (2%) and oxygen, placed them in a stereotaxic frame (Model 955, Kopf, CA) as before ^27,51^, and subcutaneously injected ketoprofen (5 mg/kg; Zoetis) for analgesia. After cleaning the rat’s head with iodide solution and applying one or two drops of lidocaine (2%, Wockhardt USA, LLC, NDC60432-464-00) to anesthetize the skin, we made a small incision to gently lift the skin and expose the skull. After making two small bilateral holes into the skulls of OTR-Cre and CRF-Cre rats, we bilaterally injected a Cre-dependent adeno-associated viral vector pAAV-hSyn-DIO-hM4D(Gi)-mCherry, which was a gift from Bryan Roth (Addgene plasmid #44362; http://n2t.net/addgene:44362; RRID:Addgene_44362) ^52^ into the BNST_DL_ (at the following coordinates from Bregma: 15° coronal angle, AP: 0.0 mm, ML: ± 3.4 mm, DV: −7.1 mm, 100 nl) using a 5-µl Hamilton syringe (Hamilton Co., Reno, Nevada) at a rate of 25 nL/min. In a separate group of male AVP-Cre rats, we injected pAAV-hSyn FLEx-mGFP-2A-synaptophysin-mRuby (gifted by Liqun Luo (Addgene plasmid #71760; http://n2t.net/addgene:71760; RRID:Addgene_71760) bilaterally into the SON (n=4, from Bregma: AP −1.44, ML +/-2.07, DV −9.047, angle 0°) or SCN (n=4, AP −0.58, ML +2.015, DV −9.1, angle 12.3°). For optogenetic experiments, we injected the SON (n=28), SCN (n=30), or PVN (n=10, coordinates from Bregma: AP −1.4, ML +/-0.6 DV −7.8 angle 0°) of another cohort of male AVP-Cre rats with a Cre-dependent AAV driving channelrhodopsin (ChR2) expression pAAV-EF1a-double floxed-hChR2(H134R)-EYFP-WPRE-HGHpA (gifted by Karl Deisseroth; Addgene plasmid #20298; http://n2t.net/addgene:20298; RRID:Addgene_20298; 100 nl, diluted 1:4 with sterile saline). A separate cohort of SD rats was injected bilaterally in the PVN (n=10) with the AAV driving ChR2-mCherry expression under the OT promoter (AAV-OTp-ChR2-mCherry, developed by Valery Grinevich ^53^). Coordinates were based on the rat brain atlas ^54^ and our prior work ^27,33,53^ and adjusted based on body weight, when needed, according to the published formula ^55^. Syringes were left in place for 8 min after viral infusion before syringe retraction. Upon completion of surgery, we sutured the skin and applied antibiotic ointment to the incision site. Rats received another injection of ketoprofen 24-hours post-surgery and were housed for another three weeks before experiments began. Fluorescence expression in brain sections from all injected transgenic rats was analyzed prior to inclusion of the animal in the analysis for electrophysiology, neuronal tract tracing, or behavioral experiments.

### Neuronal tract tracing, dual-immunofluorescence, and microscopy

Three weeks after surgeries, rats were either used for electrophysiology as above or were perfused for neuronal tract tracing. Following the electrophysiological recordings, all slices were collected, processed, and analyzed for fluorescence in the injection sites and fibers in the BNST_DL_. In the latter case, rats received an i.p. injection of euthanasia III (Covetrus, Columbus, OH) before transcardial perfusion with cold PBS (0.05 M, pH=7.4), followed by stabilized 10% formalin solution (200-250 ml, Fisher Scientific, SF98-4). Brains were dissected, stored in 10% formalin for an hour at room temperature, and then incubated in 30% sucrose-PBS for 48 h at 4°C. Frontal 50-μm-thick serial brain sections containing the BNST and the hypothalamus were sliced using a freezing-stage Microtome (model SM2000R, Leica Biosystems, Nussloch, Germany) and processed for immunofluorescence. Brain sections from AVP-Cre rats injected with a Cre-dependent-synaptophysin-mRuby into the SON or SCN for neuronal tract tracing and from AVP-Cre rats injected with a Cre-dependent-ChR2-eYFP (post-electrophysiology) were processed with AVP antibody (dilution 1:7500, rabbit, Millipore Sigma AB-1565) as before ^33^ or AVP antibody generated and kindly provided by Dr. Harold Gainer (NIH; Bethesda; USA) ^56^, and then visualized with goat anti-rabbit IgG Alexa 647 secondary antibody (dilution 1:500, Invitrogen A21245) or Alexa 594 secondary antibody (dilution 1:500, Invitrogen A11037), respectively. These sections were then mounted with Mowiol antifade reagent on gelatin-subbed slides (Fischer Scientific, 12-544-7). We first imaged mRuby or EYFP in AVP cell bodies in the SON or SCN and fibers in the BNST with a Nikon Eclipse fluorescent microscope and then used an Olympus FV10i confocal microscope (Fluoview FV10i confocal laser-scanning microscope, Olympus, Waltham, MA) to take high-magnification images and Z-stacks to visualize hypothalamic cell bodies and fibers in the BNST co-expressing mRuby or EYFP and AVP peptide.

The expression and distribution of OTR neurons in the BNST were determined on the brain sections from OTR-Cre rats (n=9) injected with a Cre-dependent AAV driving Gi-DREADD-mCherry into the BNST_DL_. To determine the phenotype of OTR neurons, we used BNST sections from these rats combined with antibodies against two enzymes involved in cellular signaling in the BNST ^35,57^, which mark mutually exclusive neuronal populations in the BNST ^58^: striatal-enriched protein tyrosine phosphatase (STEP), using anti-STEP primary antibody (mouse, dilution 1:500, Santa Cruz sc-23892) ^35^ and protein kinase C delta (PKCδ), with anti-PKCδ primary antibody (mouse, dilution 1:1000, BD Biosciences 610398) ^57^. Both proteins were then visualized with goat anti-mouse IgG Alexa 488 secondary antibody (dilution 1:500, Invitrogen A11029). Confocal microscopy was used for high-resolution images and to acquire multi-Z-stack images at 60x for cell quantification.

To count OTR-, STEP-, and PKCδ-expressing neurons and neurons co-expressing OTR and STEP or OTR and PKCδ, we took z-stacks from the entire dorsal BNST (above the anterior commissure) from 3 rats, selecting the Multi Area Z-stacks Time Lapse function in the confocal Fluoview Fv10i software. Each z-stack contained 25-35 confocal planes with 1-µm intervals with 60x magnification. Z-stacks were taken at anterior (Bregma −0.12 mm), middle (Bregma −0.24 mm), and posterior (Bregma −0.48 mm) regions of the antero-posterior axis of the BNST_DL_. OTR-, STEP-, and PKCδ-expressing neurons were counted on the left and right hemispheres from 3 sections per rat and averaged per BNST. After image acquisition, all individual z-stacks were used to create a montage of the entire dorsal BNST using the function Multi Stack Montage, which is part of the BIOP plugin package on the ImageJ 1.53t software (Image processing and analysis in Java, ^59^). The final montage was used to manually count each cell type using ImageJ’s cell counter, which allows to assign a number to each cell type and label them to prevent counting the same cell twice. OTR-mCherry and STEP or PKCδ co-expressing neurons were assigned three numbers, one for OTR-mCherry, one for STEP or PKCδ, and another to indicate double labeling. Every third consecutive BNST section (150 µm intervals) from the entire brain was used for quantification.

Diagrams and graphical abstract were created using SciDraw (images/icons created by Dr. Zhe Chen, Dr. Christophe Leterrie), Biorender (Biorender.com), Adobe Illustrator 2024, Adobe Photoshop 2024, and Microsoft PowerPoint software.

### Optogenetic stimulation protocol in AVP-Cre transgenic rats

As we previously reported detailed cellular effects of exogenous OT application in the BNST_DL_ ^40^, we first validated the optogenetic approach in the rats injected with the AAV-OTp-ChR2-mCherry virus, generated by the Grinevich Lab ^34^ in the PVN and recorded from brain slices containing the BNST. Tetanic blue light stimulation (TLS, 10-ms single pulse at 30 Hz for 20 s, which previously successfully triggered axonal OT release in the central amygdala ^53^) evoked OT release in the BNST_DL_ (**Supplementary Fig. 1**). Three weeks after Cre-dependent AAV-ChR2-eYFP was injected in the SON, PVN, or SCN, we decapitated AVP-Cre rats as above and used brain slices containing the BNST or hypothalamus for electrophysiological recordings combined with optogenetic stimulation. The slices were kept in aCSF in the dark to avoid photobleaching of the fluorescent signal and ChR2 activation. Because high frequencies of action potentials are thought to be necessary to trigger neuropeptide release, we first performed current-clamp recordings from fluorescent AVP neurons in the SON and demonstrated that 10-ms blue light pulses (BL, 470 nm) evoked action potentials in AVP SON neurons that followed frequencies up to 30 Hz applied for 20 s (**Fig. 6e**). Then, to study the effects of AVP released from terminals containing ChR2 in the BNST on the intrinsic excitability of Type I and Type III BNST_DL_ neurons, we determined the input/output (I/O) relationship in response to 80, 100, and 120pA current injections, before and after TLS. Here, we modified the optogenetic stimulation protocol to more closely mimic the physiological firing of AVP neurons in the SCN and SON ^60,61^ with 10-ms single pulses at 10 Hz for 20 s. This TLS protocol was used in BNST slices from AVP-Cre rats injected with Cre-dependent AAV-ChR2 in the hypothalamus and was delivered at the recording site using whole-field illumination through a 40x water-immersion objective (Olympus, Tokyo, Japan) with a pE-300^ultra^ CoolLED illumination system (CoolLED Ltd., Andover, UK). In a subset of neurons in each experiment, to confirm peptide receptor-mediated cellular effects, we applied the OTR antagonist, OTA (as above), before TLS.

### Acoustic startle response (ASR) and FPS in OTR-Cre rats

OTR-Cre rats (n=45) injected with a Cre-dependent AAV-Gi-DREADD-mCherry were used for chemogenetic inhibition of OTR neurons in the BNST_DL_ before fear conditioning (followed by FPS) and before EPM experiments (n=22 saline, n=21 CNO). Only rats that showed moderate to high Gi-DREADD-mCherry expression in the BNST in one or both hemispheres were considered for data analysis (n=43).

Rats were tested in Plexiglas enclosures inside sound-attenuating chambers (San Diego Instruments, Inc., CA), as described before ^27,39^. The Plexiglas enclosures were installed on top of a platform that detected movement (jump amplitude measured within a 200-ms window following the onset of the startle-eliciting noise) and transformed the velocity of the movement into a voltage output ^62^ detected by SR-Lab software (Part No: 6300-0000-Q, San Diego Instruments, Inc., CA). ASR was measured during 30 trials of startle-eliciting 95-dB white noise burst (WNB) on day 1 (chamber and startle habituation) and day 2 (baseline pre-shock test). On day 3 (fear conditioning), rats received an i.p. injection of CNO (2 mg/kg) 45 min before exposure to 10 presentations of 3.7-s cue light (conditioned stimulus, CS), each co-terminating with a 0.5-s foot shock (unconditioned stimulus, US; 0.5 mA) in context A. On day 4, rats were tested for recall of cued and non-cued fear in context B, where ASR was first measured alone (10 trials, post-shock), followed by an additional 20 trials in which ASR was measured in the presence of the cue (light+noise trials) or in the absence of the cue (noise-only trials), presented in a pseudorandom order (**Fig. 4A, Supplemental Fig.2**). Selected environmental cues were altered in context B to distinguish it from context A: context B lacked the steel grid bars used for conditioning in context A, was cleaned with a different disinfectant (ethanol 70% instead of peroxide), and had a different experimenter performing the testing. On day 5, rats were tested for contextual fear recall, where ASR was measured with no cue presentation in the original context A. To determine the rate of fear extinction, we tested rats for cued/non-cued fear three times and contextual fear two times on alternate days, see **Supplementary Fig. 2** for FPS components.

### Elevated plus maze (EPM) in OTR-Cre rats

The behavioral experiments took place in a room under a dim lighting condition. The experimental OTR-Cre rats used for the FPS test were tested on the EPM 3 days after the last FPS test. The EPM apparatus was elevated 32 inches (in) from the ground and consisted of two closed arms (width, length, height: 5 in x 20 in x 18 in), two open arms (width and length: 5 in x 20 in), and a squared center (width and length: 5 in x 5 in). Forty-five min after saline or CNO i.p. injections, we placed rats at the center of the EPM and allowed them to explore for 5 min. We measured total entries and time spent in the open arms, closed arms, and at the center using ANY-Maze 6.34 software (Stoelting, Wood Dale, IL) and quantified time spent freezing in each of these compartments.

### Functional verification of Gi-DREADD with *in vitro* whole-cell patch-clamp electrophysiology

The whole-cell patch recordings were performed as above ^40^ on 300-μm-thick brain slices containing the BNST from 5 AAV-injected OTR-Cre male rats. To confirm that OTR-BNST_DL_ neurons expressing Gi-DREADD-mCherry were functionally inhibited by CNO, we depolarized neurons to action potential threshold and recorded spontaneous activity for 5 min before bath application of CNO (20 μM), for 7 min during CNO application, and for 12 min during washout (**Fig. 4B**).

### Breeding protocols and genotyping

Rats were bred at 11-15 weeks of age by pairing Cre+/- male or female rats with wild-type SD (for OTR-Cre and AVP-Cre) or Wistar (for CRF-Cre) rats, such that all offspring were heterozygous for Cre. Wild-type breeders were purchased from Envigo, as above, from diverse litters to keep the Cre lines outbred. Pairs were together for approximately 10-14 days or until females showed signs of pregnancy and then individually housed. Rats were genotyped at weaning (postnatal day 21) for the presence of Cre. During weaning, males and females were separated and ear tagged (Braintree Scientific, Inc. 1005-1LZ). For tagging, we lightly exposed rats to the inhalant isoflurane and used an ear punch (Braintree Scientific, Inc. EP-S-902) to create a 2-mm hole for tag insertion. Ear tissue was stored in 0.2-ml 8-strip PCR tubes (GeneMate, VWR, 490003-710) labeled with the animals corresponding ID number at −20°C.

For genotyping, we added 50 μl of a lysis buffer (2.5 ml 1M Tris, pH 8.8, 100 μl 0.5M EDTApH8.0, 250ul Tween 20, 1.0 ml) with proteinase K (20 mg added to 1.0 ml 50 nM Tris-HCLpH8.0,10mM CaCl_2_ 15 μl) to each tube of tissue and placed tubes in a Thermo Cycler at 95°C overnight. The next day, we vortexed the tubes slightly and returned them to the Thermo Cycler for another 10 min at 100°C to denature the proteinase K. We then centrifuged the samples for 10 min at 6000 rpm and collected the supernatant for use with the PCR mix. The primers used were specific for the Cre-sequence within the DNA, amplifying only the DNA of Cre+ animals with the following sequence (5’ → 3’) for OTR-Cre and AVP-Cre: Cre 3 (forward) TCGCTGCATTACCGGTCGATGC (22 bp), and Cre 4 (reverse) CCATGAGTGAACGAACCTGGTCG (23 bp), and for CRF-Cre: SG13 (forward) GCATTACCGGTCGATGCAACGAGTGATGAG, and SG14 (reverse) GAGTGAACGAACCTGGTCGAAATCAGTGCG (all from ThermoFisher Scientific), as before ^41^. The PCR-mix was composed of the extracted DNA, primers, KAPA2G Fast HS Master Mix (Kapa Biosystems, Inc, KK5621), Ultrapure BSA (Invitrogen, AM2616) and nuclease free H_2_O. The PCR-Mix and protocol used for the thermal cycler is below. The PCR samples were loaded on a 2.0% agarose gel (VWR, 0710) made in 1X TAE buffer (VWR, 82021-492) with 50 μl ethidium bromide (0.625 mg/ml, VWR, E406,) in an electrophoresis chamber containing 1X TAE buffer at 80 V for 90 min. A 10-Kb DNA ladder (BioLabs, Inc., N3270S) was run in parallel to verify the size of the amplified DNA fragment.

### Statistical analysis

Electrophysiological data are presented as mean ± standard error of mean (SEM). The distribution of data sets was first evaluated with Kolmogorov-Smirnov test. When normal distribution was observed, the effects of AVP and other drugs on membrane properties were analyzed by a repeated measures (RM) one-way analysis of variance (ANOVA) or mixed-effects model, when applicable. The effects of AVP (and other pharmacological manipulations) on intrinsic excitability and firing frequency were analyzed with RM two-way ANOVA with treatment and current as factors. Where the F-ratio was significant, all-pairwise post hoc comparisons were made using Tukey’s or Sidak’s tests.

Behavioral data are presented as violin plots depicting the median and the 25^th^ and 75^th^ quartiles. FPS data were analyzed by a two-way RM ANOVA with the factors trial type (pre-shock, post-shock, noise-only, light+noise) and treatment (saline vs. CNO). When F-ratio was significant, all *post hoc* analyses were compared using Sidak’s multiple comparison test. For analyses of the effect of treatment on cued, non-cued, and contextual fear, data are presented as percentage change scores of ASR and were analyzed with un-paired *t*-tests. The percentage change of ASR was analyzed according to the following formulas:

Cued fear = (Light Noise – Noise alone) / Noise alone) x 100% in context B

Non-cued fear = (Noise alone – Post shock) / Post shock) x 100% in context B

Contextual fear = (Post shock – Pre shock) / Pre shock) x 100% in context A

Shock reactivity for each individual rat was calculated as the average startle amplitude during each of the 10 foot-shock presentations during fear conditioning (no WNB present) and was analyzed with un-paired *t*-tests, See **Supplementary Fig. 2** for the FPS components.

EPM data are presented as the total number of entries and percentage of time spent in each compartment. Total time freezing in each compartment is expressed in seconds. Comparisons between saline and CNO groups were analyzed for each parameter with an un-paired *t*-test.

All statistical analyses were completed using GraphPad Prism version 10.2.1 (339) (GraphPad Software, Inc., San Diego, CA). *P < 0.05* was considered significant.

## 3. Results

### 3.1. AVP excites Type I BNST_DL_ neurons, which requires OTR, but not V1aR or V1bR

The effects of AVP, OT, OTR and V1R antagonists and agonists on intrinsic membrane properties of Type I-III BNST_DL_ neurons are shown in **Table 1** and described in detail in the *Supplementary Results*.

**Table 1.** The effect of OTR and V1R activation on intrinsic membrane properties of Type I-III BNST_DL_ neurons.

| Type of Neuron | Treatment | Membrane Properties | Pre-Treatment<br>(Mean ± SEM) | Post-Treatment<br>(Mean ± SEM) | Statistical Analysis | p-value | n |
| --- | --- | --- | --- | --- | --- | --- | --- |
| **Type I** | **AVP** | RMP (mV) | -59.1±1.77 | -53.87±1.25 | F(2, 15)=6.340 | 0.0101 | 9 |
|  |  | Rin (MΩ) | 139.54±11.74 | 165.42±16.62 | F(1.774, 13.31)=5.695 | 0.0187 | 9 |
|  |  | Rh (pA) | 27.08±3.24 | 15.42±2.01 | F(2, 13)=7.374 | 0.0072 | 8 |
|  |  | First-spike Th (mV) | -35.83±0.79 | -35.80±0.65 | F(1.300, 9.750)=0.6786 | 0.4681 | 9 |
|  |  | First-spike Lat (ms) | 221.50±56.41 | 137.50±33.78 | F(2, 15)=7.393 | 0.0058 | 9 |
| *OTR antagonist* | **OTA + AVP** | RMP (mV) | -63.17±1.48 | -57.04±2.15 | F(1.088, 9.788)=6.851 | 0.0244 | 10 |
|  |  | Rin (MΩ) | 191.60±15.27 | 202.60±14.01 | F(1.719, 15.47)=5.789 | 0.0162 | 10 |
|  |  | Rh (pA) | 36.58±4.16 | 30.17±4.73 | F(1.584, 14.26)=1.938 | 0.1840 | 10 |
|  |  | First-spike Th (mV) | -35.38± 0.71 | -35.72±0.88 | F(1.720, 15.48)=0.3976 | 0.6484 | 10 |
|  |  | First-spike Lat (ms) | 188.20±33.61 | 134.30±32.14 | F(1.967, 17.70)=4.366 | 0.0293 | 9 |
| *V1aR antagonist* | **SR49059 + AVP** | RMP (mV) | -64.22 ± 1.93 | -62.47 ± 2.12 | F(1.571, 12.57)=5.548 | 0.0240 | 9 |
|  |  | Rh (pA) | 45.37 ± 7.00 | 38.61 ± 6.76 | F(1.806, 14.45)=6.361 | 0.0120 | 9 |
|  |  | Rin (MΩ) | 180.1 ± 15.50 | 202.1 ± 16.27 | F(1.216, 9.730)=3.353 | 0.0930 | 9 |
|  |  | First-spike Lat (ms) | 161.1 ± 18.26 | 143.4 ± 21.08 | F(1.273, 10.19)=0.7476 | 0.4393 | 9 |
|  |  | Th (mV) | -36.23 ± 1.51 | -34.88 ± 1.72 | F(1.260, 10.08)=2.523 | 0.1401 | 9 |
| *V1bR antagonist* | **Nelivaptan + AVP** | RMP (mV) | -63.13±2.28 | -56.76±2.73 | F(1.352, 12.17)=15.50 | 0.0010 | 10 |
|  |  | Rin (MΩ) | 189.5±16.30 | 214.6±17.92 | F(1.421, 12.79)=4.299 | 0.0480 | 10 |
|  |  | Rh (pA) | 39.50±5.70 | 31.00±5.36 | F(1.387, 12.48)=8.143 | 0.0093 | 10 |
|  |  | First-spike Th (mV) | -30.28±1.69 | -28.59±2.22 | F(1.292, 11.63)=2.541 | 0.1335 | 10 |

| Type of Neuron | Treatment | Membrane Properties | Pre-Treatment<br>(Mean ± SEM) | Post-Treatment<br>(Mean ± SEM) | Statistical Analysis | p-value | n |
| --- | --- | --- | --- | --- | --- | --- | --- |
| | | First-spike Lat (ms) | 187.2±28.40 | 113.6±18.15 | F(1.707, 15.37)=6.963 | 0.0090 | 10 |
| **Type II** | **AVP** | RMP (mV) | -53.12±1.46 | -51.29±1.69 | F(1.155, 6.927)=2.219 | 0.1815 | 8 |
| | | Rin (MΩ) | 135.9±15.83 | 151.3±21.57 | F(2, 12)= 3.348 | 0.0699 | 8 |
| | | Rh (pA) | 25.10±5.39 | 25.62±8.73 | F(1.016, 6.095)=0.08300 | 0.7867 | 8 |
| | | First-spike Th (mV) | -36.82±0.82 | -34.65±1.53 | F(1.818, 10.91)=1.199 | 0.3336 | 8 |
| | | First-spike Lat (ms) | 97.3±7.3 | 75.27±8.7 | F(1.403, 8.415)=1.146 | 0.3397 | 8 |
| **Type III** | **AVP** | RMP (mV) | -68.34±1.36 | -65±1.93 | F(2, 15)=5.717 | 0.0143 | 10 |
| | | Rin (MΩ) | 108.8±10.29 | 128.9±13.83 | F(1.602, 5.608)=30.90 | 0.0011 | 5 |
| | | Rh (pA) | 75.15±9.45 | 63.5±9.65 | F(1.114, 8.356)=12.18 | 0.0068 | 10 |
| | | First-spike Th (mV) | -33.12±1.89 | -33.46±1.92 | F(1.187, 8.902)=0.4559 | 0.5495 | 10 |
| | | First-spike Lat (ms) | 419.08±64.09 | 257.5±52.13 | F(1.869, 13.08)=11.79 | 0.0014 | 10 |
| | **OTR/V1aR Antagonist + AVP** | RMP (mV) | -69.79±1.21 | -67.97±1.17 | F(1.211, 7.264)=3.616 | 0.0936 | 7 |
| | | Rin (MΩ) | 94.74±16.92 | 97.37±17.56 | F(1.904, 11.42)=2.037 | 0.1761 | 7 |
| | | Rh (pA) | 88.33±8.97 | 83.33±4.42 | F(1.335, 8.007)=0.8376 | 0.4213 | 7 |
| | | First-spike Th (mV) | -33.41±0.95 | -34.69±0.79 | F(1.344, 8.063)=2.478 | 0.1517 | 7 |
| | | First-spike Lat (ms) | 203.2±29.08 | 176.2±32.81 | F(1.174, 7.041)=0.3909 | 0.5843 | 7 |
| | **OTA + AVP** | RMP (mV) | -67.37±1.72 | -66.59±1.9 | F(1.039, 6.235)=0.6159 | 0.4675 | 7 |
| | | Rin (MΩ) | 110.99±8.5 | 132.79±15. | F(1.042, 4.169)=5.504 | 0.0758 | 5 |
| | | Rh (pA) | 77.02±10.75 | 75.71±12.74 | F(1.197, 7.185)=0.0635 | 0.8495 | 7 |
| | | First-spike Th (mV) | -38.86±1.96 | -37.58±2.46 | F(1.926, 11.56)=1.482 | 0.2667 | 7 |
| | | First-spike Lat (ms) | 327.6±60.45 | 300.3±60.50 | F(2, 10)=0.5830 | 0.5761 | 6 |
| | **SR49059 + AVP** | RMP (mV) | -66.95 ± 1.88 | -63.94 ± 2.38 | F(1.341, 9.386)=11.51 | 0.0052 | 8 |
| | | Rin (MΩ) | 159 ± 13.93 | 181.5 ± 14.85 | F(1.764, 12.35)=5.944 | 0.0179 | 8 |
| | | Rh (pA) | 68.96± 7.48 | 56.67 ± 7.13 | F(1.249, 8.745)=7.536 | 0.0193 | 8 |
| | | First-spike Th (mV) | -34.09 ± 1.33 | -33.25 ± 2.33 | F(1.844, 12.91)=1.825 | 0.2014 | 8 |
| | | First-spike Lat (ms) | 404 ± 72.91 | 261.2 ± 65.96 | F(1.546, 10.83)=7.551 | 0.0121 | 8 |
| | **Nelivaptan + AVP** | RMP (mV) | -74.24 ± 1.46 | -69.45 ± 1.74 | F(1.331 10.65) = 15.67 | 0.0014 | 9 |
| | | Rin (MΩ) | 124.9 ± 9.76 | 186.6 ± 24.8 | F(1.196, 9.565) = 9.060 | 0.0114 | 9 |
| | | Rh (pA) | 94.44 ± 5.86 | 84.26 ± 6.5 | F(1.507, 12.06) = 5.441 | 0.0272 | 9 |
| | | First-spike Th (mV) | -29.9 ± 1.76 | -24.24 ± 3.12 | F(1.077, 8.613) = 4.558 | 0.0610 | 9 |
| | | First-spike Lat (ms) | 377.2 ± 64.25 | 248.4 ± 37.86 | F(1.539, 12.31) = 4.491 | 0.0402 | 9 |
| **Type III** | **OT** | RMP (mV) | -68.54±1.03 | -67.79±0.87 | F(2, 13)=1.374 | 0.2875 | 8 |
| | | Rin (MΩ) | 137.15±23.61 | 142.41±26.16 | F(0.8062, 3.628)=14.14 | 0.0251 | 6 |
| | | Rh (pA) | 83.02±4.81 | 69.33±4.74 | F(2, 13)=13.22 | 0.0007 | 8 |
| | | First-spike Th (mV) | -29.27±1.86 | -29.49±1.98 | F(0.04181, 0.2717)=2.479 | 0.1480 | 8 |
| | | First-spike Lat (ms) | 497.8±63.95 | 281.05±43.37 | F(1.827, 10.96)=11.13 | 0.0027 | 8 |
| *OTR agonist* | **TGOT** | RMP (mV) | -71.86±1.00 | -70.00±1.22 | F(1.168, 8.177)=5.241 | 0.0469 | 9 |
| | | Rin (MΩ) | 85.79±6.63 | 91.24±7.02 | F(1.076, 4.841)=8.456 | 0.0338 | 7 |
| | | Rh (pA) | 93.61±9.88 | 82.69±10.97 | F(1.776, 11.54)=6.129 | 0.0174 | 9 |
| | | First-spike Th (mV) | -30.84±0.98 | -30.79±1.13 | F(1.120, 7.838)=0.6998 | 0.4439 | 9 |
| | | First-spike Lat (ms) | 594.03±76.46 | 355.24±68.69 | F(0.9019, 5.412)=7.726 | 0.0373 | 9 |
| *V1aR agonist* | **FE 201874** | RMP (mV) | -70.72±0.9 | -69.44±1.25 | F(1.242, 9.312)=3.093 | 0.1068 | 10 |
| | | Rin (MΩ) | 118.21±1.25 | 117.57±9.69 | F(1.219, 6.094)=0.2128 | 0.70748 | 7 |
| | | Rh (pA) | 77.83±9.55 | 73.66±10.5 | F(1.209, 9.066)=1.728 | 0.2256 | 10 |
| | | First-spike Th (mV) | -30.45±1.22 | -31.93±1.1 | F(1.550, 11.63)=2.515 | 0.1312 | 10 |
| | | First-spike Lat (ms) | 432.8±65.72 | 345.3±57.68 | F(1.710, 11.12)=3.780 | 0.0610 | 9 |
| *V1bR agonist* | **d[Cha4]-AVP** | RMP (mV) | -70.42±0.84 | -69.81±1.15 | F(1.987, 12.91)=1.327 | 0.2988 | 8 |
| | | Rin (MΩ) | 105.18±7.85 | 110.98±10.05 | F(0.9314, 6.054)=2.663 | 0.1532 | 8 |
| | | Rh (pA) | 86.25±5.96 | 81.46±6.2 | F(1.160, 7.540)=1.193 | 0.3195 | 8 |
| | | First-spike Th (mV) | -33.38±1.67 | -34.00±1.79 | F(0.9257, 6.017)=2.467 | 0.1666 | 8 |
| | | First-spike Lat (ms) | 475.1±59.48 | 426.6±60.20 | F(1.063, 5.848)=10.24 | 0.0185 | 8 |

#### 3.1.1. The effect of AVP on intrinsic excitability of Type I BNST_DL_ neurons

We measured steady state firing frequency (SSF) as a function of current (input/output, I/O, relationship) before (pre), during AVP application, and after (washout, post). There was a significant incremental current effect on the action potential frequency (*P*=0.0002, F(1.267,10.14)=29.52) and a significant treatment effect, with AVP inducing a leftward shift of the I/O relationship, without affecting its slope (*P*=0.0137, F(1.201, 9.611)=8.415). There was no interaction between current and treatment (*P*=0.9013, F (1.862,11.55)=0.09175, n=9, mixed-effect analysis). *Post hoc* test showed a significant AVP effect on SSF at (in pA) 50 (*P*=0.0298), 60 (*P*=0.0274), 70 (*P*=0.0255), and 80 (*P*=0.0181, vs. pre), **Fig.1A-A’**. When the OTR antagonist, OTA, was applied before AVP, there was a current effect on spike frequency (*P*<0.0001, F(1.341, 10.73)=59.21, n=9, mixed-effects model) but no treatment effect (*P*=0.0574, F(1.466, 11.73)=3.999) and no interaction (*P*=0.1642, F(1.688, 8.440)=2.281), **Fig.1B-B’**. In the presence of the V1aR antagonist, SR49059, there was a current effect on spike frequency (*P*<0.0001, F(1.312, 10.49)=52.74, n=9), and AVP in a presence of SR49059 significantly changed the I/O relationship (*P=*0.0002, F(1.886, 15.09)=16.29), with no interaction (*P*=0.1137, F(1.708, 7.075)=3.065). Multiple comparisons showed an excitatory effect of AVP in the presence of SR49059 (vs. pre) at all currents injected from 40 to 180pA (40 (*P*=0.0355), 50 (*P*=0.0180), 60 (*P*=0.0189), 70 (*P*=0.0009), 80 (*P*=0.0014), 90 (*P*=0.0092), 100 (*P*=0.0035), 110 (*P*=0.0032), 120 (*P*=0.0070), 130 (*P*=0.0111), 140 (*P*=0.0069), 150 (*P*=0.0052), 160 (*P*=0.0060), 170 (*P*=0.0067), and 180 (*P*=0.0021)). Notably, although SR49059 did not block the AVP effect, it showed an excitatory effect on its own vs. pre at (pA) 70 (*P*=0.0323), 80 (*P*=0.0221), 90 (*P*=0.0370), 100 (*P*=0.0328), 110 (*P*=0.0328), 120 (*P*=0.0376), 150 (*P*=0.0397), 160 (*P*=0.0432), 170 (*P*=0.0493), and 180 (*P*=0.0374), **Fig.1C-C’**. When the V1bR antagonist, Nelivaptan, was applied before AVP, there was a current effect on spike frequency (*P*<0.0001, F(2.294, 20.65)=117.9, n=10). However, AVP in the presence of Nelivaptan significantly shifted the I/O relationship (*P*<0.0001, F(1.651, 14.86)=42.15), without interacting with the current (*P*=0.4980, F(1.027, 5.475)=0.5382). Multiple comparisons showed a significant excitatory effect of AVP on SSF at (in pA) 30 (*P*=0.0175), 40 (*P*=0.0282), 50 (*P*=0.0184), 60 (*P*=0.0042), 70 (*P*=0.0017), 80 (*P*=0.0034), 90 (*P*=0.0050), 100 (*P*=0.0155), 110 (*P*=0.0159), 120 (*P*=0.0048), 130 (*P*=0.0103), 140 (*P*=0.0083), 150 (*P*=0.0268), and 160 (*P*=0.0256) but not at 170 (*P*=0.2845) or 180 (*P*=0.2847), **Fig.1D-D’**.

**Figure 1.**
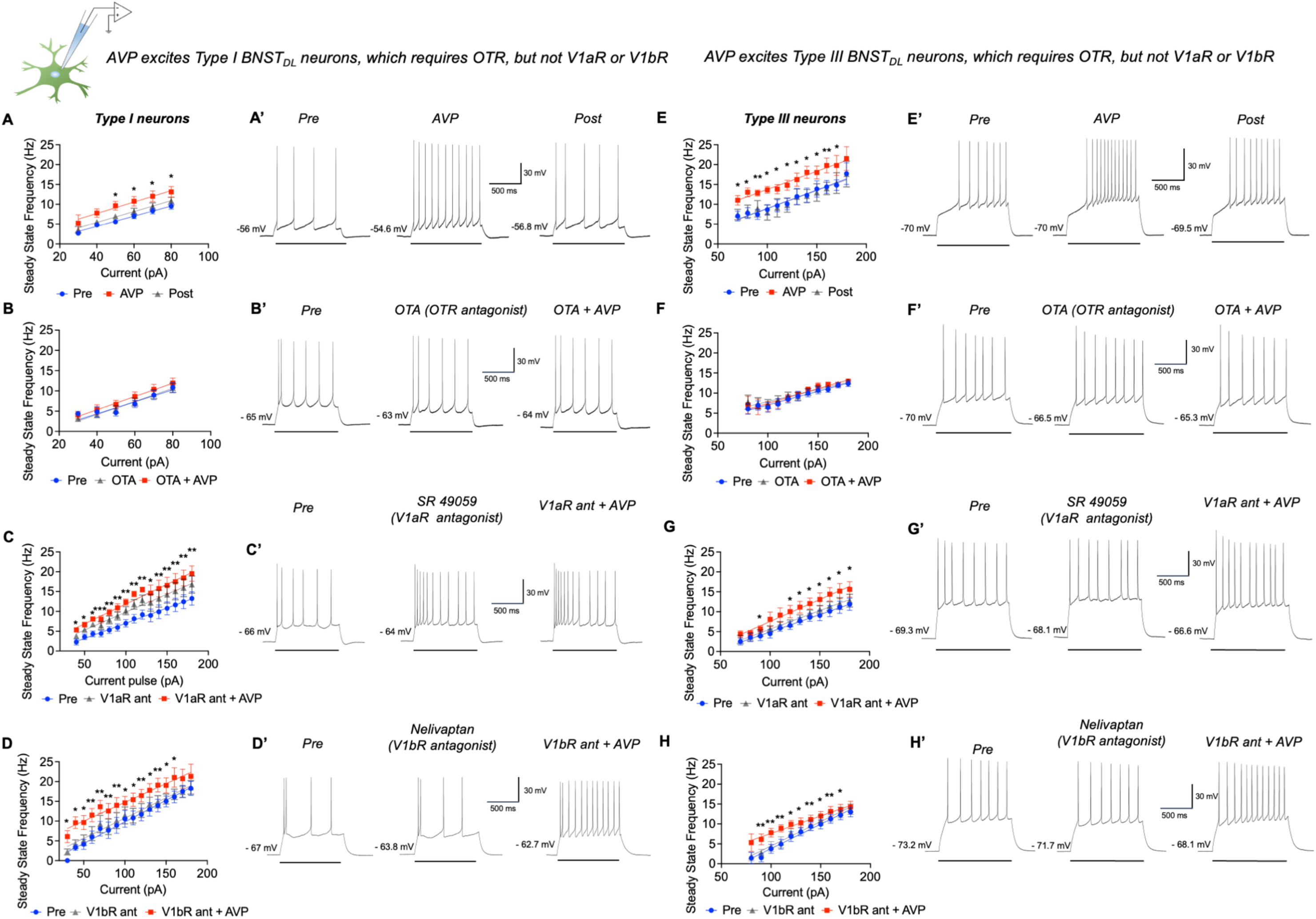
Arginine-vasopressin (AVP) increases intrinsic excitability of Type I and Type III BNST_DL_ neurons, which requires OTR, but not V1aR or V1bR transmission. **A)** <u>In Type I neurons</u>, AVP induced a leftward shift of the I/O relationship, without affecting its slope (*P*=0.0137, n=11, mixed-effect analysis). The change in SSF induced by AVP completely recovered during washout for all currents injected (in pA): 50 (*P*=0.1187), 60 (*P*=0.2704), 70 (*P*=0.2469), and 80 (*P*=0.2121, pre vs. post). **A’)** Representative traces of Type I BNST neuron responses to a depolarizing current pulse of 40pA for 1 sec before (Pre), during (AVP) and after AVP (Post). **B)** Application of OTR antagonist, OTA, blocked the excitatory effect of AVP in Type I neurons (*P*=0.0574, n=9). **B’)** Representative traces of Type I neuron responses to a depolarizing current pulse of 60pA for 1 sec in the presence of OTA. **C)** In the presence of the V1aR antagonist, SR49059, AVP application significantly changed the I/O relationship (*P=*0.0002, n=9) and multiple comparisons showed an excitatory effect of AVP at all currents injected. **C’)** Representative traces of Type I neuron responses to a depolarizing current pulse of 100pA for 1 sec in the presence of SR49059. **D)** In a presence of V1bR antagonist, Nelivaptan, AVP significantly shifted the I/O relationship (*P*<0.0001, n=10). Multiple comparisons showed an excitatory effect of AVP on SSF at all current injected except 170 (*P*=0.2845) or 180pA (*P*=0.2847). Nelivaptan alone had no effect on the SSF at any of the currents injected except at 140pA (*P*=0.0474). **D’)** Representative traces of Type I neuron responses to a depolarizing current pulse of 80pA for 1 sec during Nelivaptan. **E)** <u>In Type III neurons</u>, AVP induced a leftward shift of the I/O relationship, without affecting its slope (*P*=0.0007, n=11), with a significant AVP effect in all currents injected except 180 (*P*=0.0584). The leftward shift of the I/O relationship in Type III neurons recovered during washout of AVP for all currents injected**. E’)** Representative traces of Type III neuron responses to a depolarizing current pulse of 140pA before, during (AVP) and after AVP (Post). **F)** in the presence of OTR antagonist, OTA, there was no treatment effect of AVP on the I/O relationship (*P*=0.3949, n=9). **F’)** Representative traces of Type III neuron response to a depolarizing current pulse of 100pA during OTA. **G)** In the presence of the V1aR antagonist, SR49059, AVP application significantly changed the I/O relationship (*P*=0.0068, n=7). Multiple comparisons showed an excitatory effect of AVP on SSF at most current injected except (in pA) at 70 (*P*=0.1359), 80 (*P*=0.2954), 100 (*P*=0.1874), and 110 (*P*=0.1720). Application of SR49059 alone did not have an effect on the SSF at any of the currents injected. **G’)** Representative traces of Type III neuron response to a depolarizing current pulse of 140pA during SR49059. **H)** When the V1bR antagonist, Nelivaptan, was applied before AVP, AVP significantly shifted the I/O relationship (*P*<0.0001, n=10). Multiple comparisons showed a significant excitatory effect of AVP on SSF at most currents injected except 80 (*P*=0.1881) and 180pA (*P*=0.0578). **H’)** Representative traces of Type III neuron responses to a depolarizing current pulse of 110pA during Nelivaptan. * *P*<0.05, ** *P*<0.01, *** *P*<0.001 vs. pre

### 3.2. AVP does not affect intrinsic excitability of Type II BNST_DL_ neurons

Mixed-effect analysis showed a significant incremental effect of currents injected on spike frequency in Type II BNST_DL_ neurons (*P*<0.0001, F(1.256, 8.789)=79.04, n=8). However, AVP did not affect SSF (*P*=0.1749, F(1.173, 8.210)=2.211), and there was no interaction between current and treatment (*P*=0.3589, F(1.414, 8.200)=1.074, n=8), **not shown**.

### 3.3. AVP excites Type III BNST_DL_ neurons, which requires OTR, but not V1aR or V1bR

#### 3.3.1. The effect of AVP on intrinsic excitability of Type III BNST_DL_ neurons

There was a significant incremental current effect on action potential frequency (*P*=0.0036, F(0.5547, 5.547)=29.81, n=11, mixed-effect analysis), while AVP also induced a leftward shift of the I/O relationship, without affecting its slope (*P*=0.0007, F(0.9966, 9.966)=23.05), with no interaction between current and treatment (*P=*0.3127, F(1.669,5.463)=1.410), *Post-hoc* analysis showed a significant AVP effect (in pA) at 70 (*P*=0.0209), 80 (*P*=0.0283), 90 (*P*=0.0076), 100 (*P*=0.0116), 110 (*P*=0.0319), 120 (*P*=0.0226), 130 (*P*= 0.0453), 140 (*P*=0.0174), 150 (*P*=0.0487), 160 (*P*=0.0099), and 170 (*P*=0.0486, pre vs. AVP) and trending at 180 (*P*=0.0584). The leftward shift of the I/O relationship in Type III neurons recovered during washout of AVP for all currents injected (*P*=0.2824-0.9845, pre vs. post), **Fig.1E-E’**. In the presence of the OTR antagonist, OTA, there was a significant incremental current effect (*P*<0.0001, F(1.877, 15.01)=61.34, n=9) but no treatment effect of AVP on the I/O relationship (*P*=0.3949, F(1.419, 11.35)=0.9149, n=9), and no interaction (*P*=0.3986, F(0.6373, 1.880)=0.8231), suggesting that blocking OTR completely abolished the excitatory effects of AVP in Type III neurons, **Fig.1F-F’**. In contrast, in the presence of the V1aR antagonist, SR49059, there was a significant current effect on spike frequency (*P*=0.0003, F(0.9437, 5.662)=62.36, n=7), and AVP application significantly changed the I/O relationship (*P*=0.0184, F(1.166, 6.997)=8.869), with no interaction (*P*=0.3410, F(1.224, 4.841)=1.202). Application of SR49059 alone did not have an effect on the SSF of Type III neurons at any of the currents injected (*P*>0.05), **Fig.1G-G’**. When the V1bR antagonist, Nelivaptan, was applied before AVP, there was a significant current effect on spike frequency (*P*<0.0001, F(2.294, 20.65)=5117.9, n=10, mixed-effect analysis). However, AVP in the presence of Nelivaptan significantly shifted the I/O relationship (*P*<0.0001, F(1.651, 14.86)=42.15, n=10), with no interaction with the current (*P*=0.4980, F(1.027, 5.475)=0.5382, n=10), **Fig.1H-H’**. Application of AVP in the presence of the V1aR/OTR antagonist showed a significant increase in spike frequency as the current increased (*P*<0.0001, F(1.543, 10.80)=55.46, n=8, mixed-effect analysis); however, AVP did not change the I/O relationship (*P*=0.2742, F(0.6225, 4.358)=1.293), **Fig.2A**.

**Figure 2.**
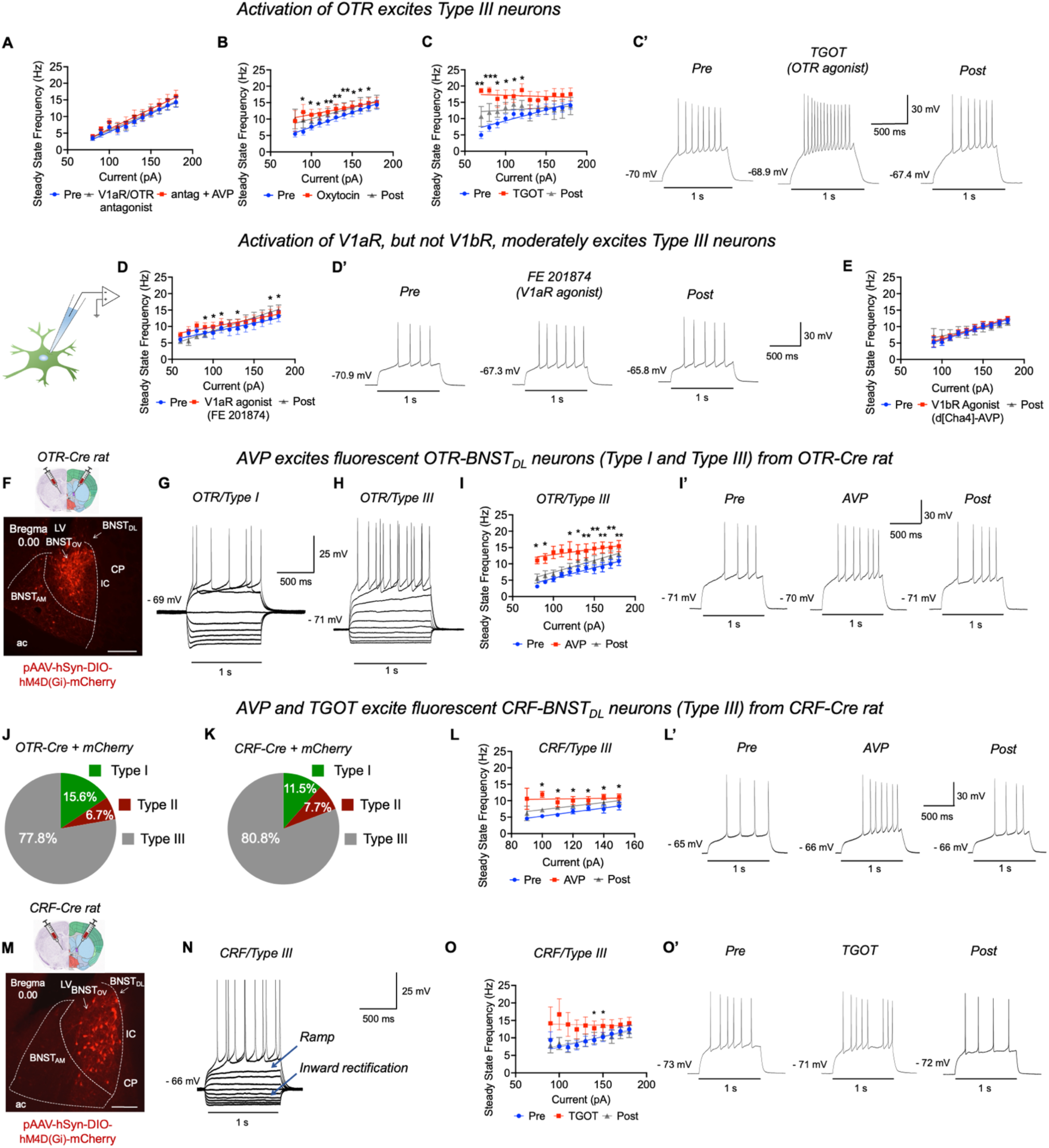
Activation of OTR robustly excites OTR and CRF-expressing Type III BNST_DL_ neurons, whereas activation of V1aR, but not V1bR, has a modest excitatory effect. **A)** In Type III neurons from wild-type rats, application of AVP in the presence of the V1aR/OTR antagonist ((d(CH_2_)_5_^1^,Tyr(Me)^2^,Arg^8^)-vasopressin) did not change the I/O relationship (*P*=0.2742, n=8). **B)** Application of endogenous OTR agonist oxytocin (OT) induced a leftward shift of the I/O relationship (*P*=0.0077, n=8), with significant OT effect on SSF in most currents injected except at 80 (*P*=0.3822) and 180pA (*P*=0.2500). **C)** Selective and potent OTR agonist, TGOT, affected the I/O relationship (*P*=0.0196, n=8) with significant excitatory effect from 70pA to 110pA and trending at 120 (*P*=0.0673). These effects were fully reversed with washout at (in pA) 70 (*P*=0.1571), 80 (*P*=0.1040), 90 (*P*=0.1552), 100 (*P*=0.1417), and 110 (*P*=0.1115, pre vs. post). **C’)** Examples of evoked responses of a Type III neuron to a depolarizing pulse of 90pA during TGOT application. **D)** Selective V1aR agonist, FE201874, induced an excitatory effect on the I/O relationship (*P*=0.0117, n=10), with *post hoc* analysis showing significant effects at (in pA): 90 (*P*=0.0451), 100 (*P*=0.0392), 110 (*P*=0.0287), 130 *P*=0.0455), 170 (*P*=0.0381), and 180 (*P*=0.0327), which were fully reversed at washout. **D’)** Examples of evoked responses of a Type III neuron to a depolarizing pulse of 90pA for 1 sec during FE201874 application. **E)** Selective V1bR agonist, d[Cha4]-AVP, did not induce a significant effect on I/O relationship (*P*=0.5334, n=8), * *P*<0.05, ** *P*<0.01, *** *P*<0.001 vs. pre. **F)** Nissl (left) and anatomical annotations (right) from the Allen Mouse Brain Atlas and Allen Reference Atlas - Mouse Brain ^114^. High somatodendritic OTR-mCherry expression in the BNST_DL_ after male OTR-Cre rats were injected with a Cre-dependent AAV-DREADDs-mCherry in the BNST_DL_. **G)** Example of evoked responses of a Type I/OTR mCherry fluorescent neuron and **H)** Type III/OTR mCherry neuron to a hyperpolarizing and depolarizing current pulses of 1 sec. **I)** In Type III/OTR fluorescent neurons, mixed-effect analysis revealed a significant incremental effect of AVP on SSF (*P*<0.0001, n=10), with a significant excitatory effect of AVP on SSF at all current injected except trends at 100pA (*P*=0.0678) and 110pA (*P*=0.0763). The effect of AVP in Type III/OTR neurons was reversed during the washout of AVP at most currents injected except at (pA) 130 (*P*=0.0445), 150 (*P*=0.0500), 160 (*P*=0.0450), 170 (*P*=0.0445), and 180 (*P*=0.0294). **I’)** Representative trace of Type III/OTR fluorescent neuron responses to a depolarizing current pulse of 130pA for 1 sec during AVP. **J)** Fluorescent OTR-BNST_DL_-mCherry neurons were characterized as Type I-III neurons based on their electrophysiological properties, with majority 35/45 (77.8%) characterized as Type III, 7/45 (15.6%) as Type I, and 3/45 (6.7%) as Type II neurons. **K)** The great majority of recorded fluorescent CRF-BNST_DL_-mCherry neurons (21/26) were characterized as Type III (80.8%), whereas 2/26 as Type II (7.7%), and 3/26 as Type I (11.5%). **L)** In Type III/CRF neurons from CRF-Cre transgenic rats, there was a significant incremental effect of AVP on SSF (*P*=0.0057, n=7), with multiple comparisons showing significant effect of AVP at all current injected except at 90pA (*P*=0.2288). **L’)** Representative traces of Type III/CRF fluorescent neuron responses to a depolarizing current pulse of 100pA for 1 sec during AVP. **M)** Nissl (left) and anatomical annotations (right) from the Allen Mouse Brain Atlas and Allen Reference Atlas - Mouse Brain ^114^. High somatodendritic CRF-mCherry expression in the BNST_DL_ after male CRF-Cre transgenic rats were injected with a Cre-dependent AAV-DREADDs-mCherry in the BNST_DL_. **N)** Example of evoked responses of a Type III/CRF fluorescent neuron to a hyperpolarizing and depolarizing current pulses of 1 sec. **O)** In Type III/CRF neurons, there was a significant TGOT effect on action potential frequency (*P*=0.0167, n=7). Multiple comparisons showed a significant effect of TGOT at (in pA): 140 and 150, and trends at 130 (*P*=0.0807), 160 (*P*=0.0978), and 170 (*P*=0.0867). **O’)** Representative traces of Type III/CRF fluorescent neuron responses to a depolarizing current pulse of 150pA during TGOT application. * *P*<0.05, ** *P*<0.01, *** *P*<0.001 vs pre, ac – anterior commissure, BNST_AM_ – anteromedial BNST, BNST_OV_ – oval nucleus of the BNST, CP – Caudate Putamen, IC – Internal Capsulae, LV - Lateral Ventricle.

#### 3.3.2. The effect of OTR activation on intrinsic excitability of Type III BNST_DL_ neurons

OT induced a significant incremental current effect on spike frequency (*P*<0.0001, F(1.323,9.261)=86.87, n=8, mixed-effect analysis), and a leftward shift of the I/O relationship (*P*=0.0077, F(1.826, 12.79)=7.565), with no interaction between current and treatment (*P*=0.1023, F(1.004, 4.921)=4.015). *Post-hoc* analysis showed a significant OT effect (in pA) at 90 (*P*=0.0436), 100 (*P*=0.0218), 110 (*P*= 0.0392), 120 (*P*=0.0026), 130 (*P*= 0.0043), 140 (*P*=0.0088), 150 (*P*=0.0284), 160 (*P*=0.0204), and 170 (*P*=0.0449). The effects of OT were not significant at (in pA) 80 (*P*= 0.3822) or 180 (*P*= 0.2500), **Fig.2B**. In the presence of OTR agonist, TGOT, there was an incremental current effect on action potential frequency (*P*=0.0003, F(1.421, 9.944)=24.05, n=8), and a significant effect on I/O relationship (*P*=0.0196, F(0.8587, 6.011)=10.48) with no interaction (*P*=0.1344, F(2.149, 4.004)=3.443). TGOT had a significant excitatory effect at (in pA) 70 (*P*=0.0022), 80 (*P*=0.0001), 90 (*P*=0.0350), 100 (*P*=0.0272), and 110 (*P*=0.0394, pre vs. TGOT) and trending at 120 (*P*=0.0673), **Fig.2C-C’**.

#### 3.3.3. The effect of V1aR and V1bR activation on intrinsic excitability of Type III BNST_DL_ neurons

When FE201874 was tested as a specific agonist of V1aR, mixed-effect analysis revealed a significant effect of current injected on spike frequency (*P*<0.0001, F(1.809, 16.28)=60.13, n=10), and a significant effect on the I/O relationship (*P*=0.0117, F(1.283, 11.54)=8.023), with no interaction between current and treatment (*P*=0.1567, F(1.967, 6.230)=2.535). Multiple comparison showed a significant effect of FE201874 on SSF at (in pA) 90 (*P*=0.0451), 100 (*P*=0.0392) 110 (*P*=0.0287), 130 (*P*=0.0455), 170 (*P*=0.0381), and 180 (*P*=0.0327) and trends at 140 (*P*=0.0618) and 150 (*P*=0.0684). These effects of FE201874 were reversed at all of these currents injected (*P*=0.8557, *P*=0.9721, *P*=0.1264, *P*=0.2114, *P*=0.1975, *P*=0.1807, respectively, pre vs. post), **Fig.2D-D’**. In the presence of V1bR agonist, d[Cha4]-AVP, there was an incremental current effect on spike frequency (*P*<0.0001, F(1.123, 7.859)=34.12, n=8, mixed-effect analysis), with no significant effect on I/O relationship (*P*=0.5334, F(0.2668, 1.867)=0.02568), and no interaction (*P*=0.2199, F(0.4503, 1.551)=2.697), **Fig.2E**.

### 3.4. AVP and TGOT excite Type III/CRF and Type III/OTR neurons from transgenic rats

#### 3.4.1. AVP excites Type III/OTR neurons from OTR-Cre rats

Intrinsic excitability of fluorescent OTR neurons was measured as SSF in slices obtained from male OTR-Cre rats injected with Cre-dependent AAV driving mCherry expression in the BNST_DL_. First, fluorescent OTR-mCherry neurons were characterized as Type I-III neurons based on their electrophysiological properties. We recorded from a total of 45 OTR-mCherry neurons, from which 7 (15.6%) were characterized as Type I, 3 (6.7%) as Type II, and 35 (77.8%) as Type III neurons (**Fig.2G-H, 2J**). In Type III/OTR fluorescent neurons, mixed-effect analysis revealed a significant current effect on spike frequency (*P*<0.0001, F(0.9525, 8.573)=59.68, n=10). Notably, AVP showed a significant incremental effect on SSF (*P*<0.0001, F(1.267,11.40)=31.89, mixed-effect analysis), with a significant interaction between current and treatment (*P=*0.0168, F(2.795, 9.364)=5.850). Multiple comparisons showed a significant incremental effect of AVP respective to baseline (pre) at (in pA) 80 (*P*=0.0130) and 90 (*P*=0.0251), trends at 100 (*P*=0.0678) and 110 (*P*=0.0763), and a significant effect at 120 (*P*=0.0240), 130 (*P*=0.0140), 140 (*P*=0.0077), 150 (*P*=0.0040), 160 (*P*=0.0032), 170 (*P*=0.0036), and 180 (*P*=0.0044). The effect of AVP in Type III/OTR neurons was reversed during the washout of AVP at (in pA) 80 (*P*=0.2158, pre vs. post), 90 (*P*=0.2818), 100 (*P*=0.1373), 110 (*P*=0.3799), 120 (*P*=0.1994), and 140 (*P*=0.0863), as multiple comparison showed no significant difference from baseline, but was not reversed at 130 (*P*=0.0445), 150 (*P*=0.0500), 160 (*P*=0.0450), 170 (*P*=0.0445), or 180 (*P*=0.0294), **Fig.2I-I’**.

#### 3.4.2. AVP excites Type III/CRF neurons from CRF-Cre rats

Previous studies demonstrated that the majority of electrophysiologically defined Type III neurons of the BNST_DL_ express mRNA for CRF at a single-cell level ^35,36^. Therefore, we tested the effect of AVP on the intrinsic excitability, measured as SSF, of fluorescent CRF neurons recorded in slices obtained from male CRF-Cre rats injected with a Cre-dependent AAV driving mCherry expression in the BNST_DL_. The great majority of recorded fluorescent CRF-mCherry neurons (n=26) were characterized as Type III (n=21, 80.8%), whereas n=2 were classified as Type II (7.7%), and n=3 as Type I neurons (11.5%), **Fig.2K, 2N**. In Type III/CRF neurons, there was a significant effect of current injected on spike frequency (*P*=0.0475, F(1.216, 7.293)=5.387, n=7), and AVP showed a significant incremental effect on SSF (*P*=0.0057, F(1.392, 8.351)=11.82, n=7, mixed-effect analysis), with a trend in the interaction between current and treatment (*P=*0.0573, F(1.511, 3.399)=7.655), **Fig.2L-L’**.

#### 3.4.3. TGOT excites CRF/Type III neurons from CRF-Cre rats

In Type III/CRF neurons, there was a significant effect of current injected on spike frequency (*P*=0.0003, F(1.723, 10.34)=21.78, n=7, mixed-effect analysis), and TGOT induced a significant incremental effect on SSF (*P*=0.0167, F(1.647, 9.882)=6.823), with a trend in the interaction between current and treatment (*P=*0.0583, F(1.669, 6.585)=4.686), **Fig.2O-O’**.

### 3.5. The BNST_DL_ contains numerous OTR neurons, which co-express STEP or PKCδ

A Numerous OTR-mCherry cell bodies and spiny dendrites were localized in the anterior (bregma 0.20 to −0.00 mm, **Fig.3b-c**), middle (bregma −0.26 to −0.35 mm, **Fig.3d-e**), and posterior (bregma - 0.60 mm, **Fig.3f**) subdivisions of the BNST. The majority of the OTR neurons were located in the BNST_DL_, primarily clustered in the oval nucleus of the BNST_DL_ (BNSTov, Bregma 0.00 mm to −0.35 mm, **Fig.3c-e**), whereas fewer OTR neurons were found in the anteromedial BNST (BNST_AM_, **Fig.3b-e**). In posterior BNST sections, OTR neurons were found in lateral division, posterior BNST (BNST_LP_), lateral division, intermediate posterior BNST (BNST_LI_), and to a lesser extent in medial, posterolateral BNST (BNST_MPL_ **Fig.3f**). Double-immunofluorescence analysis of confocal images from the BNST_DL_ showed that 16.91±4.41% of OTR-mCherry neurons co-expressed PKCδ (**Fig.3k**), whereas only 7.87±1.08% of all PKCδ neurons co-expressed OTR-mCherry (**Fig.3l**). Double-immunofluorescence toward STEP showed that 18.23±2.05% of all OTR-mCherry neurons co-expressed STEP (**Fig.3p**), whereas 20.22±2.51% of all STEP neurons co-expressed OTR-mCherry (**Fig.3r**). These results suggest that while OTR neurons constitute a diverse population of BNST_DL_ neurons, STEP is the most reliable marker of OTR-BNST_DL_ neurons identified so far.

**Figure 3.**
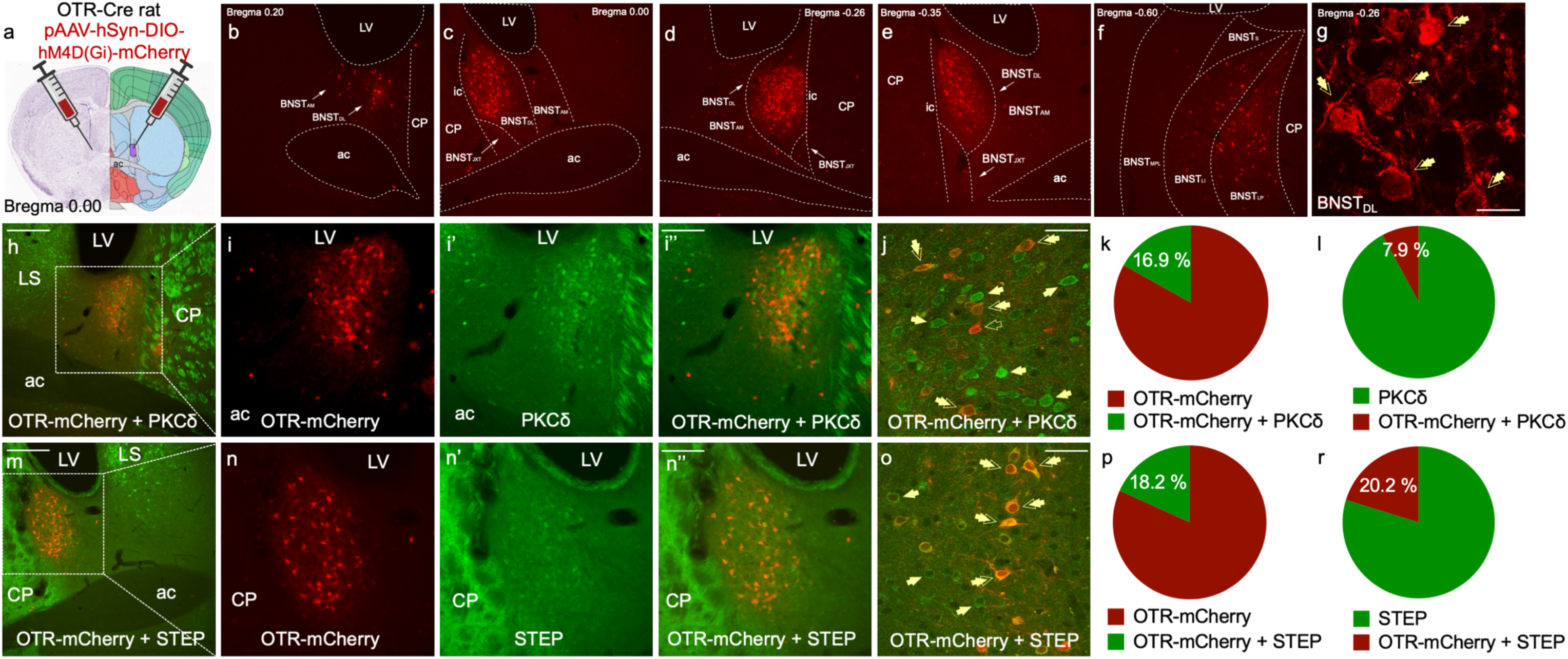
The BNST_DL_ contains numerous OTR-expressing neurons, which co-express Striatal-enriched Protein Tyrosine Phosphatase (STEP) or Protein Kinase C delta (PKCδ). Nissl (left) and anatomical annotations (right) from the Allen Mouse Brain Atlas and Allen Reference Atlas – Mouse Brain ^114^ **(a)**. OTR-mCherry cell bodies and spiny dendrites are localized in the anterior (**b-c**), middle (**d-e**), and posterior (**f**) subdivisions of the BNST. The majority of the OTR neurons were located in the BNST_DL_, primarily in the oval nucleus of the BNST_DL_ (BNSTov, **c-e**), whereas fewer OTR neurons are seen in the anteromedial BNST (BNST_AM_). In posterior BNST sections (**f)**, OTR neurons are found in lateral division, posterior BNST (BNST_LP_), lateral division, intermediate posterior BNST (BNST_LI_), and to a lesser extent in medial, posterolateral BNST (BNST_MPL_). High magnification confocal image shows robust somatodendritic expression of mCherry in OTR-BNST_DL_ neurons (scale bar 10 μm) (**g**). Double-immunofluorescence microphotographs show high density of OTR and PKCδ neurons in the BNST_DL_ (10x, **h**) and co-expression between the two populations (20x, **i-i’’**). Double-immunofluorescence confocal images (**j,** 60x, bar 20 μm) and quantification of neurons from multi-tiles Z-stacks of the entire BNST_DL_ shows that 16.91±4.41% of OTR-mCherry neurons co-express PKCδ (**k**), whereas only 7.87±1.08% of all PKCδ neurons co-express OTR-mCherry (**l**). High density of OTR and STEP neurons is found in the BNST_DL_ (10x, **m**) with high co-expression between the two populations (20x, **n-n’’**). Double-immunofluorescence confocal images (**o,** 60x, bar 20 μm) and quantification of neurons from multi-tiles Z-stacks of the entire BNST_DL_ show that 18.23±2.05% of all OTR-mCherry neurons co-express STEP (**p**), whereas 20.22±2.51% of all STEP neurons co-express OTR-mCherry (**r**).

### 3.6. Activity of OTR-BNST_DL_ neurons attenuates threat-elicited vigilant state measured in the FPS

The experimental timeline of behavioral experiments is shown in **Fig.4A**. First, the inhibitory effect of DREADDs-Gi in OTR-BNST_DL_ neurons was confirmed with patch-clamp electrophysiology recordings in brain slices containing the BNST from OTR-Cre rats (n=4) injected with a Cre-dependent AAV-DREADDs-Gi-mCherry. Here, application of CNO (20 µM) significantly reduced the spontaneous firing rate recorded near the threshold for action potentials (**Fig.4B**).

**Figure 4.**
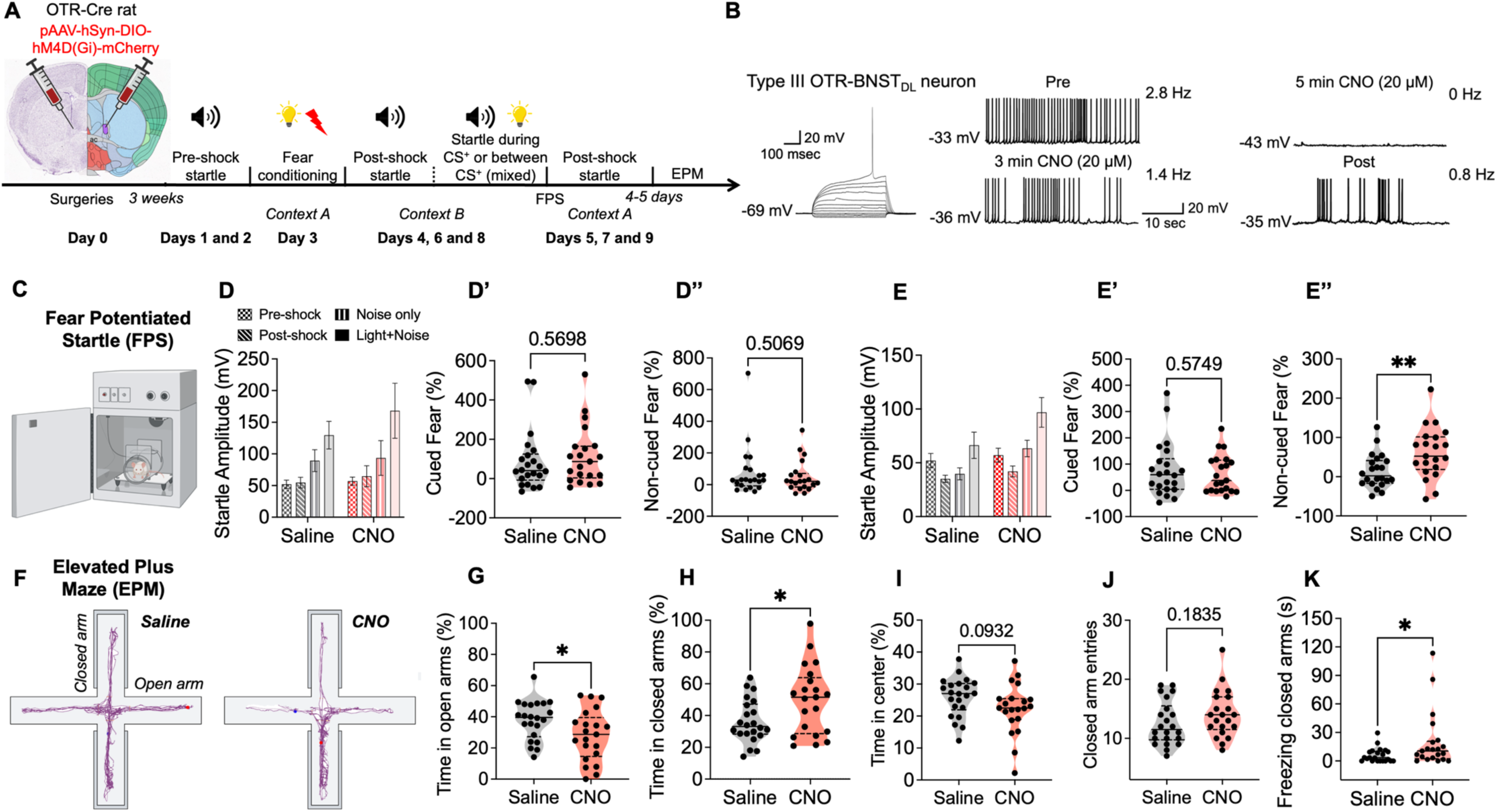
Activity of OTR-neurons in the BNST_DL_ reduces expression of anxious arousal measured in the FPS and increases open arms exploration in the EPM. **A)** Nissl (left) and anatomical annotations (right) from the Allen Mouse Brain Atlas and Allen Reference Atlas – Mouse Brain ^114^. Experimental timeline shows that male OTR-Cre rats were injected with a AAV-DIO-DREADDs-Gi-mCherry into the BNST_DL_ and were treated with CNO or saline before fear-conditioning in the FPS and once again, before the EPM. **B)** In vitro whole-cell patch-clamp electrophysiology in brain slices from the OTR-Cre rats shows that CNO (bath applied continuously for 7 minutes) reduces frequency of spontaneous firing of fluorescent OTR-BNST-mCherry neurons, which recovers during 12 minutes washout (post). **C**) FPS apparatus with Plexiglass animal enclosure and startle detection system underneath the stage. **D-D’’)** During the first FPS recall, two-way RM ANOVA showed a significant trial effect of noise only vs. light+noise (*P*<0.0001), and post-shock vs. noise only trials (*P*=0.0109), but no significant treatment effects (**D**). No significant differences were found in the percentage change of cued fear (**D**’), and non-cued fear (**D’’**). **E-E’’)** During the second FPS recall, there was a significant trial effect between the noise-only and light+noise trials (*P*<0.0001) and a significant treatment effect (*P*=0.0477). There was also a significant trial effect between the post-shock and noise-only trials (*P*=0.0024), a significant treatment effect (*P*=0.0307), and interaction (*P*=0.0430, **E**). *Post-hoc* analysis showed that CNO-treated rats (*P*=0.0012) but not saline-treated rats (*P*=0.6559), exhibited a significantly higher startle in the noise-only trials than in the post-shock trials. There were no differences in the percentage change of cued fear (**E’**) but the percentage change of non-cued fear (cue-induced vigilant state or anxious arousal) was significantly higher in CNO-treated rats than in saline controls (*P*=0.0077, **E’’**). **F)** Representative traces of locomotor activity during the EPM exploration by OTR-Cre rats treated with saline or CNO. Rats treated with CNO spent significantly less time in the open arms of the EPM (**G**), spent more time in the closed arms (**H**), and tended to spend less time in the center (**I**) vs. saline controls. No significant differences were found between rats treated with saline vs. CNO in the number of entries to the closed arms (**J**). CNO-treated rats spend significantly more time freezing in the closed arms vs. saline-treated rats (*P*=0.0344). * *P*<0.05, ** *P*<0.01

OTR-Cre rats (n=43) injected with the AAV-DREADDs as above were used to assess the effect of chemogenetic inhibition of OTR-BNST_DL_ neurons before fear conditioning. All OTR-Cre rats included in the behavioral analysis showed high to moderate expression of mCherry in OTR-BNST_DL_ neurons in one or both hemispheres. During fear conditioning, there were no significant differences in the reactivity to foot-shocks between saline- and CNO-injected rats (*P*=0.1041, saline n=22, CNO n=21, **not shown**). During the first FPS recall in context B, there was a significant trial effect between the noise-only and light+noise trials (*P*=0.0001, F(1, 41)=18.25, two-way RM ANOVA), indicating effective cued fear learning, but no significant effect of treatment (saline vs. CNO) (*P*=0.5831, F(1, 41)=0.3060), nor significant interaction between trial and treatment (*P*=0.2086, F(1, 41)=1.632, **Fig.4D**). There was also a significant trial effect between the post-shock and noise-only trials (*P*=0.0109, F(1, 41)=7.121), indicating a non-cued fear expression, but no significant treatment effect (*P*=0.7613, F(1, 41)=0.09352), nor significant interaction (*P*=0.8035, F(1, 41)=0.06275, **Fig.4D**). During the first contextual fear recall in context A, there was a significant trial effect between the pre-shock and post-shock trials (*P*=0.0459, F(1, 41)=4.239) but no significant treatment effect (*P*=0.8054, F(1, 41)=0.06150), nor interaction (*P*=0.2545, F(1, 41)=1.336, **not shown**). During the first FPS recall test, no significant differences were found in the percentage change of cued fear (*P*=0.5698, **Fig.4D’**), non-cued fear (*P*=0.5069, **Fig.4D’’)**, or contextual fear (*P*=0.2541, **not shown**).

During the second FPS recall in context B, there was a significant trial effect between the noise-only and light+noise trials (*P*<0.0001, F(1, 41)=24.68) and a significant treatment effect (saline vs. CNO) (*P*=0.0477, F(1, 41)=4.168) but no significant interaction (*P*=0.5685, F(1, 41)=0.3305, **Fig.4E**). *Post-hoc* analysis showed that light+noise startle tended to be higher in the CNO group than in the saline group (*P*=0.0768). There was also a significant trial effect between the post-shock and noise-only trials (*P*=0.0024, F(1, 41)=10.52), a significant treatment effect (*P*=0.0307, F(1, 41)=5.009), and a significant interaction between trials and treatment (*P*=0.0430, F(1, 41)=4.361). *Post-hoc* analysis showed that CNO-treated rats (*P*=0.0012) but not saline-treated rats (*P*=0.6559, **Fig.4E**) exhibited a significantly higher ASR in the noise-only trials than in the post-shock trials. During the second contextual fear recall test in context A, two-way RM ANOVA showed no significant trial effect between the pre-shock and post-shock trials (*P*=0.9590, F(1, 41)=0.002675), no significant treatment effect (*P*=0.7974, F(1, 41)=0.06678), and no interaction (*P*=0.5209, F(1, 41)=0.4193, **not shown**). There were no significant differences in the percentage change of cued fear (*P*=0.5749, **Fig.4E’**) or contextual fear (p=0.6199, **not shown**) in the second FPS test. However, there was a significantly higher percentage change of non-cued fear in CNO-treated rats than in saline-treated rats (*P*=0.0077, **Fig.4E’’**). These results show that OTR-BNST_DL_ neuron activity reduces long-term expression of non-cued fear, and hence attenuates cue-induced vigilant state or anxious arousal.

### 3.7. Activity of OTR-BNST_DL_ neurons increases exploration of open arms in the EPM without affecting locomotor activity

The same OTR-Cre rats (n=43) injected with a Cre-dependent AAV-Gi-DREADD-mCherry into the BNST_DL_ were next used to examine the effect of chemogenetic inhibition of OTR-BNST_DL_ neurons on exploratory behavior in the EPM. Examples of the locomotor activity of saline- vs. CNO-treated rat are shown in **Fig.4F**. No significant differences between saline and CNO-treated rats were found in the number of entries to the open arms (*P*=0.3382, unpaired *t*-test, **not shown**), closed arms (*P*=0.1835, **Fig.4J**), or the center of the EPM (*P*=0.8387, **not shown**), suggesting that silencing OTR-BNST_DL_ neurons does not affect locomotor activity in the EPM. Time spent in each compartment of the EPM was calculated as the percentage of total time in the EPM. CNO-treated OTR-Cre rats spent significantly less time in the open arms (*P*=0.0307, **Fig.4G**) and significantly more time in the closed arms (*P*=0.0168, **Fig.4H**) and tended to spend less time at the center (*P*=0.0932, **Fig.4I**) than saline-treated controls. No significant differences were found in time freezing in open arms (*P*=0.3066, **not shown**), but CNO-treated rats spent significantly more time freezing in closed arms (*P*=0.0344, **Fig.4K**) than saline-treated rats.

### 3.8. Hypothalamic AVP neurons from the SON, SCN, and the PVN project to the BNST_DL_

Male AVP-Cre rats injected bilaterally into the SON, SCN, or PVN with a Cre-dependent pAAV-hSyn-FLEx mGFP-2A-synaptophysin-mRuby were perfused (n=8) or injected with pAAV-EF1a-double floxed-hChR2(H134R)-eYFP-WPRE-HGHpA, used for slice electrophysiology and examined *post-hoc* for neuronal projections from the hypothalamic AVP neurons to the BNST_DL_ (SON n=28; SCN n=30; PVN n=10). Fidelity of the Cre expression in the hypothalamic AVP neurons was confirmed in the cell bodies and processes of AVP neurons in the SON, SCN, and PVN, where synaptophysin-mRuby (**Fig.5a-b’’’,5d-e’’’**) or EF1a-ChR2-eYFP-expressing neurons and processes co-expressed AVP peptide (**Fig.6a-d’’**,**7a-f’’’**). Fibers and processes expressing mRuby or eYFP were seen to ascend from the SCN neurons and traversed dorsally along the 3^rd^ ventricle, innervating the anterior hypothalamus and the PVN, including the PVN AVP neurons (**Fig.7a’’’’**), and then continued via a right angle toward the posterior BNST and dorsally toward the lateral septum (**Fig.7b’’**). From the SON, mRuby or eYFP-expressing fibers and processes ascended via a lateral-dorsal route toward the hypothalamic accessory nuclei (AN) and the PVN (**Fig.6a-c’’**), and then laterally along the internal capsule (IC) toward the posterior BNST (**Fig.6g**). From the PVN, eYFP-expressing fibers and processes descended toward SON, and travelled via a lateral route toward the internal capsule (**Fig.7e**). The eYFP-expressing fibers and processes originating from the SON (**Fig.6b-c’’**), SCN (**Fig.7b’-b’’**), and PVN (**Fig.7f’’’**) co-expressed mature AVP peptide. Notably, both synaptophysin-mRuby and EF1a-ChR2-eYFP-expressing fibers originating from the SON (**Fig.5f-f’’’,6f-f’’**), SCN (**Fig.5c-c’’’,7c-c’’’**), and PVN (**Fig.7g-g’’’**), were found in the BNST_DL_, and a subset of these fibers co-expressed mature AVP peptide.

**Figure 5.**
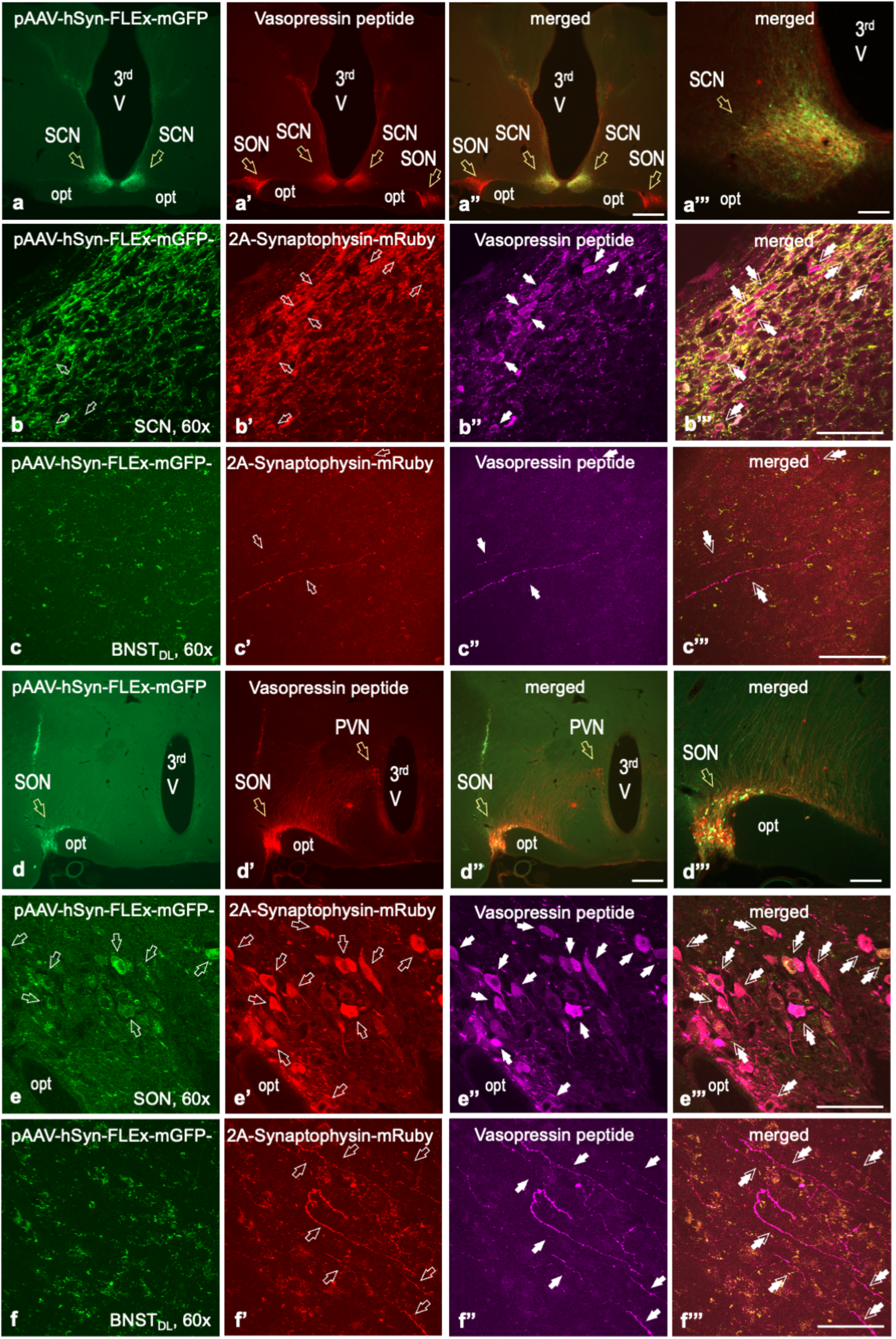
Suprachiasmatic nucleus and supraoptic nucleus of the hypothalamus send AVP-containing peptidergic fibers to the BNST_DL_. AVP-Cre transgenic rats were bilaterally injected with pAAV-hSyn-FLEx-mGFP-2A-Synaptophysin-mRuby in the SCN or SON and AVP peptide was visualized by Alexa Flour 647. **a-a’’’**) Co-expression of GFP (green) and AVP peptide (red) in the SCN, but no SON contamination with GFP, is observed at 4x (**a’’**, scale bar 100 µm) and 20x (**a’’’**, bar 25 µm) magnification. **b-b’’’**) Confocal images (60x, bar 10 µm) show the co-expression of hSyn-GFP (green, **b**), 2A-Synaptophysin-mRuby (red, open arrows, **b’**), and AVP peptide (magenta, closed arrows, **b’’**) in neurons of the SCN (double arrows, **b’’’**). **c-c’’’**) Confocal images (60x) showing the co-localization of 2A-Synaptophysin-mRuby (red, open arrows, **c’**) and AVP peptide (magenta, closed arrows, **c’’**) in fibers of the BNST_DL_ that originated from the SCN (scale bar 10 µm, double arrows, **c’’’**). **d-d’’’**) Co-expression of GFP (green) and AVP peptide (red) in the SON, but no SCN contamination with GFP, is observed at 4x (**d’’**, bar 100 µm) and 20x (**d’’’**, 25 µm) magnification. **e-e’’’**) Confocal images (60x, bar 10 µm) show co-expression of hSyn-GFP (green, **e**), 2A-Synaptophysin-mRuby (red, open arrows, **e’**), and AVP peptide (magenta, closed arrows, **e’’**) in neurons of the SON (double arrows, **e’’’**). **f-f’’’**) Confocal images (60x, bar 10 µm) show co-expression of 2A-Synaptophysin-mRuby (red, open arrows, **f’**) and AVP peptide (magenta, closed arrows, **f’’**) in fibers of the BNST_DL_ that originated from the SON (double arrows, **f’’’**), opt – optic track, 3^rd^V – 3^rd^ Ventricle.

**Figure 6.**
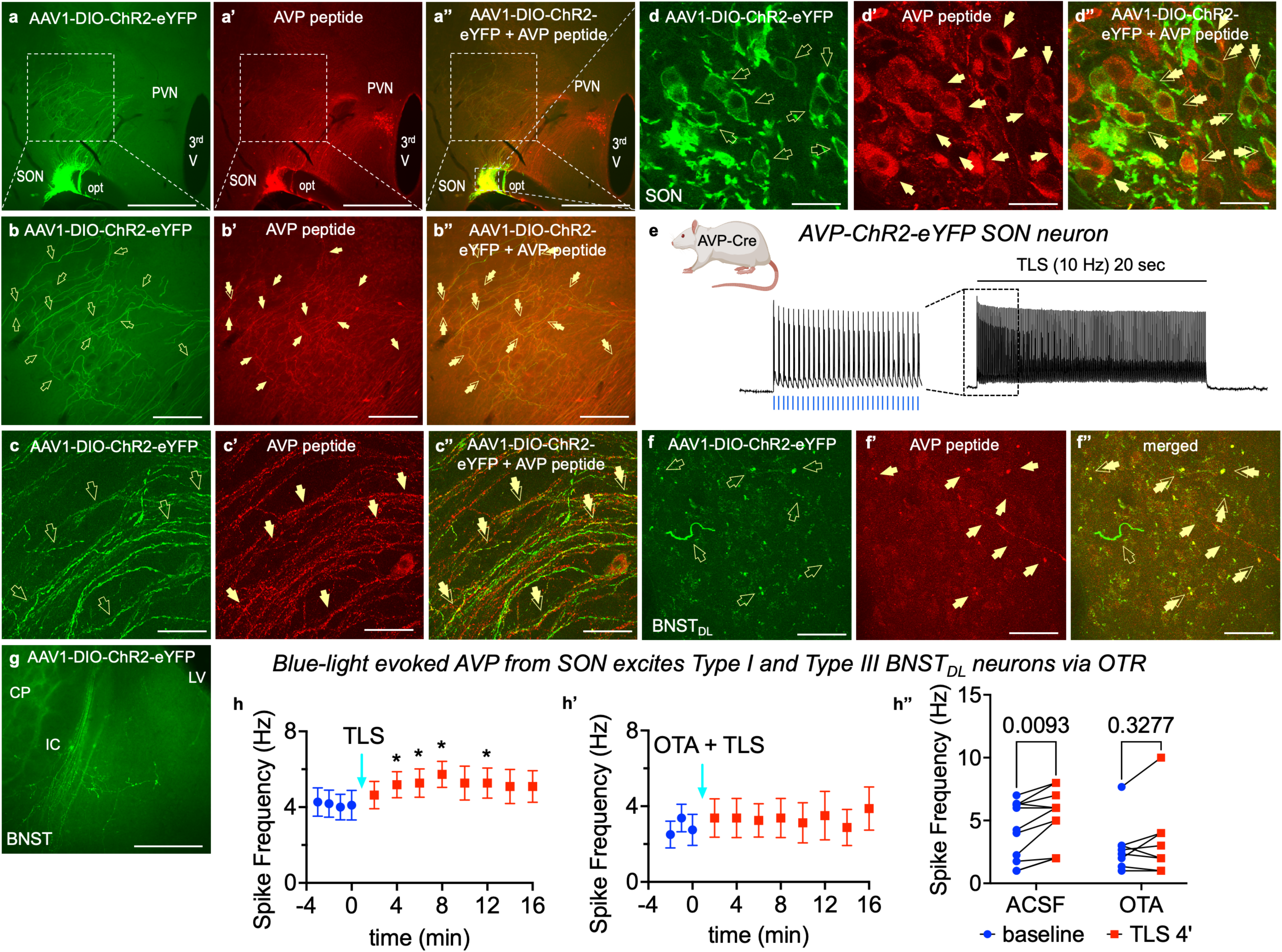
Tetanic light stimulation of AVP fibers from the SON excite BNST_DL_ neurons in an OTR-dependent fashion. AVP-Cre transgenic rats were bilaterally injected with AAV-EF1a-DIO-ChR2-eYFP (green) in the SON and AVP peptide was visualized with Alexa Flour 594 (red). Co-expression of eYFP-ChR2 and AVP peptide in the SON, but not the PVN, is observed at 4x magnification (**a-a’’**, scale bar 100 μm). High co-expression of ChR2-eYFP (green, open arrows) and AVP peptide (red, closed arrows) is seen in SON-originating fibers, which ascend dorso-laterally from the SON toward internal capsulae (IC), as seen at 20x (**b-b’’**, bar 50 μm) and 60x magnification (**c-c’’** confocal images, bar 10 μm). Confocal images (60x) show high co-localization of ChR2-eYFP (green, open arrows) and AVP (red, closed arrows) in the soma and dendrites of SON neurons (**d-d’’**, bar 10 μm). In the BNST_DL_, confocal images show ChR2-eYFP-expressing (green, open arrows) and AVP-containing fibers (red, closed arrows) converging on boutons which co-express ChR2-eYFP and AVP peptide (double arrows, bar 10 μm, **f-f’’**). Action potentials recorded from fluorescent ChR2-eYFP-expressing SON neurons from brain slices containing hypothalamus show burst firing following tetanic blue light stimulation (TLS, 10 Hz) for 20 sec, where each pulse of blue light evokes action potential (**e**). High expression of fluorescent ChR2-eYFP-expressing processes and fibers with fluorescent boutons (presumptive peptide release sites), originating from the SON, is seen in the BNST from brain slice processed *post-electrophysiology* (scale bar 100 μm, **g**). In brain slices containing the BNST, action potentials frequency (Hz) was recorded from identified Type I and Type III BNST_DL_ neurons, before and after TLS (TLS, 10 Hz) of hypothalamic ChR2-eYFP fibers in response to 80pA current injection, with and without bath application of the OTR antagonist, OTA. In Type I and Type III BNST_DL_ neurons, analyzed together, there was a significant TLS effect on firing frequency (*P*=0.0459, n=11, **h**) but not in the presence of OTR antagonist, OTA (*P*=0.2863, n=8, **h’**). Comparison of firing frequency with and without OTA at baseline vs. 4 min, in response to 80pA current, for all neurons recorded showed a significant TLS effect (*P*=0.0055), with increased firing following TLS, but not in the presence of OTA. CP – Caudate Putamen, LV – Lateral Ventricle, 3^rd^V – 3^rd^ Ventricle, * *P*<0.05

**Figure 7.**
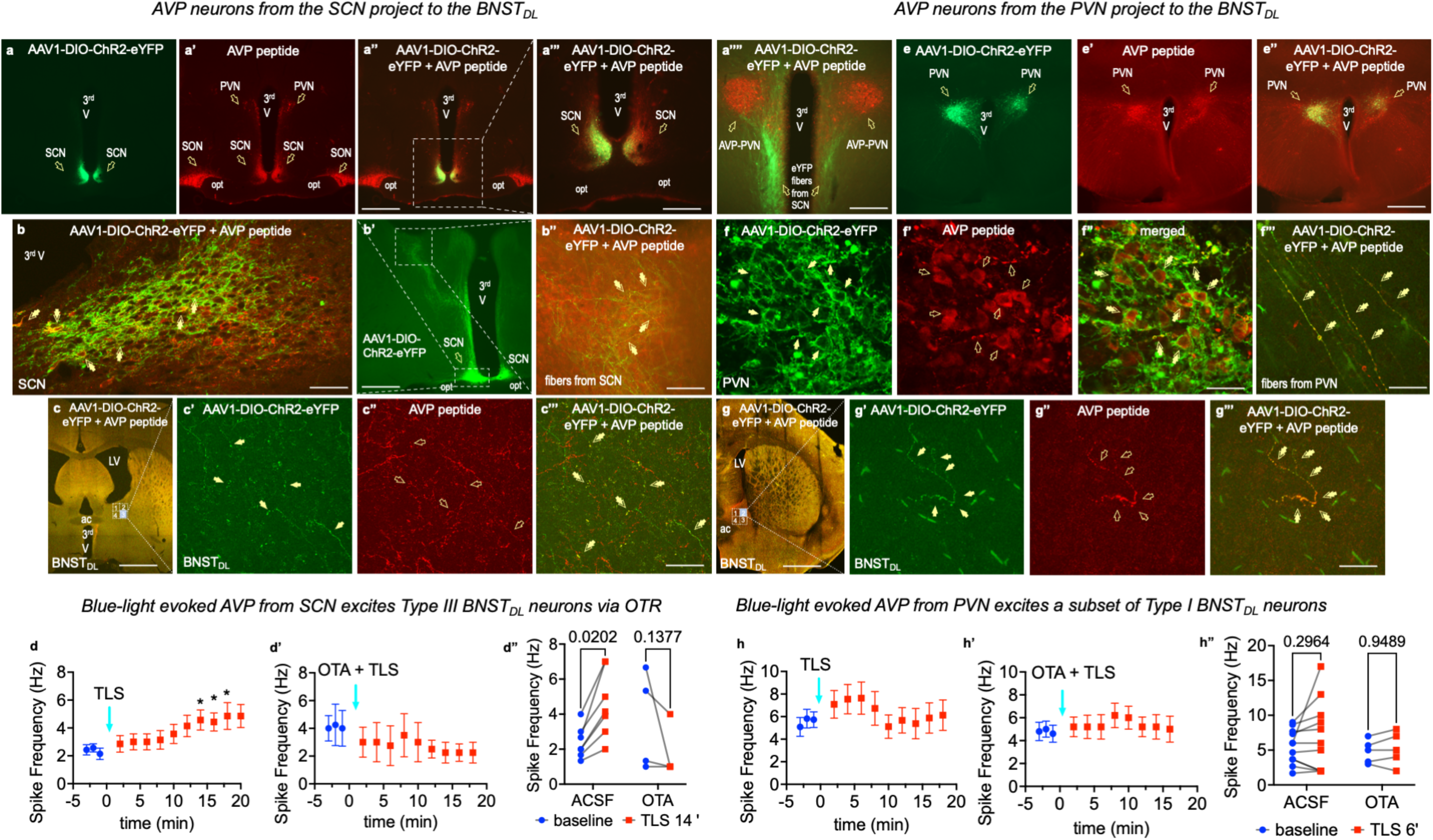
Tetanic light stimulation of AVP fibers from the SCN or PVN excites BNST_DL_ neurons in an OTR-dependent fashion. AVP-Cre transgenic rats were bilaterally injected with AAV-EF1a-DIO-ChR2-eYFP (green) in the SCN (**a-b’’**) or PVN (**e-g’’’**) and AVP peptide was visualized with Alexa Flour 594 (red). *<u>In rats injected in the SCN</u>*, co-expression of eYFP-ChR2 and AVP peptide in the SCN, but not SON or PVN, is observed at 4x (**a-a’’**, scale bar 100 μm) and 10x magnification (**a’’’**, bar 50 μm). High co-expression of ChR2-eYFP (green) and AVP peptide (red) is seen in the SCN neurons (**b**, double arrows, confocal image, 60x, bar 10 μm) and SCN-originating fibers, which ascend dorsally from the SCN toward the PVN (**a’’’’**, **b’**, 10x), and travel at a straight angle toward posterior BNST (**b’**, 10x), where they co-express AVP peptide (**b’’**, double arrows, 20x, bar 20 μm). In the BNST_DL_, confocal images show ChR2-eYFP-expressing (green, open arrows) and AVP-containing fibers and processes (red, closed arrows) which co-express ChR2-eYFP and AVP peptide (double arrows, 60x, scale bar 10 μm, **c-c’’’**). In brain slices containing the BNST, action potentials frequency (Hz) was recorded from identified Type I and Type III BNST_DL_ neurons, before and after TLS (10 Hz) of SCN-originating ChR2-eYFP fibers in response to 80pA current injection, with and without OTR antagonist, OTA. In Type III BNST_DL_ neurons, there was a significant TLS effect (*P*=0.0251, n=7, RM ANOVA), with multiple comparisons showing significant increases in firing frequency at 14, 16, and 18 min following TLS (**d**). In the presence of the OTR antagonist, OTA, TLS had no effect on firing of Type III neurons (*P*=0.4261, n=4, **d’**). When firing following TLS was analyzed with and without OTA, 2-way ANOVA showed a significant interaction between the TLS and OTA treatment (*P*=0.0059, n=11) with increased firing frequency following TLS but not in the presence of OTA (**d’’**). *<u>In rats</u> <u>injected in the PVN</u>*, co-expression of eYFP-ChR2 and AVP peptide is observed in the PVN, but not SON (**e-e’’**, 4x, scale bar 100 μm). High co-expression of ChR2-eYFP (closed arrows, **f**) and AVP peptide (open arrows, **f**’) is seen in the PVN neurons (double arrows, **f’’**, confocal image, 60x, bar 10 μm) and PVN-originating fibers (**f’’’’**, 60x). In the BNST_DL_, confocal images show ChR2-eYFP-expressing (**g’**, green, closed arrows) and AVP-containing fibers and processes (**g’’**, red, closed arrows) which co-express ChR2-eYFP and AVP peptide (double arrows, 60x, bar 10 μm, **g’’’**). In brain slices containing the BNST, action potentials frequency (Hz) was recorded from identified Type I and Type III BNST_DL_ neurons, before and after TLS (10 Hz) of PVN-originating ChR2-eYFP fibers in response to 80pA current injection, with and without OTR antagonist, OTA. In Type I and III BNST_DL_ neurons, there was no TLS effect (*P*=0.3404, n=11, **h**) and no TLS effect in a presence of OTA (*P*=0.2920, n=5, **h’**). No treatment or TLS effect (*P*=0.3028, n=16) emerged when firing following TLS was analyzed with and without OTA (**h’’**). However, individual Type I BNST_DL_ neurons (2/7), from rats with a very high bilateral PVN ChR2-eYFP expression, showed significant spiking increase to 283 and 144% of baseline at 6 min post TLS, whereas no Type I neurons showed increase in firing frequency when OTA was applied before TLS (**h’’**). ac – anterior commissure, CP – Caudate Putamen, LV – Lateral Ventricle, opt – optic track, 3^rd^V – 3^rd^ Ventricle, * *P*<0.05

### 3.9. Optogenetically-evoked release of AVP from hypothalamic fibers excites BNST_DL_ neurons

To determine functional peptidergic projections between the hypothalamic nuclei producing AVP and BNST_DL_ neurons, action potentials frequency (Hz) was recorded from Type I and Type III BNST_DL_ neurons, before and after tetanic light stimulation (TLS, 10 Hz) of hypothalamic ChR2-eYFP fibers in BNST slices. These recordings were performed in the presence of synaptic transmission blockers in response to 80, 100, and 120-pA current injections to determine if optogenetically released AVP mimics the excitatory effects of exogenous AVP described above. Here, we stimulated ChR2-eYFP fibers originating from the SON, SCN, and PVN, separately, with and without bath application of the OTR antagonist, OTA. At least 3-4 pre-TLS responses to each current injection were collected for every neuron and averaged as a baseline for repeated measures analyses. Only rats with strong bilateral, strong unilateral, or moderate bilateral ChR2-eYFP expression levels in the hypothalamus were included in the analyses below.

#### 3.9.1 AVP from SON fibers excites Type I and Type III BNST_DL_ neurons via OTR

First, we demonstrated that TLS (10-ms single pulses at 10 Hz for 20 s) evoked action potentials of fluorescent eYFP-ChR2 expressing AVP neurons in the SON at each light pulse delivered (**Fig.6d-e**). Following TLS of ChR2-expressing fibers in the BNST_DL_, in Type I neurons, there was a significant current effect on spiking frequency (*P*=0.0283, F(1.234, 7.404)=6.912, n=7, 2-way RM ANOVA) but no TLS effect (*P*=0.5131, F(1.630, 9.777)=0.6497) and no interaction between current and TLS (*P*=0.3164, F(1.918, 11.51)=1.267). Similarly, in Type III neurons, there was a significant current effect on firing (*P*=0.0123, F(0.5618, 2.247)=58.26, n=4, 2-way RM ANOVA) but no TLS effect (*P*=0.2023, F(1.405, 5.621)=2.137), nor interaction (*P*=0.5631, F(1.782, 6.147)=0.5937).

When the Type I and Type III neurons were pulled together, there was no significant overall TLS effect (*P*=0.1212, F(2.804, 28.04)=2.138), n=11,) despite a current effect (*P*<0.0004, F(1.204, 12.04)=21.51), but when the responses to 80 pA were analyzed separately, there was a significant TLS effect (*P*=0.0459, F(3.166, 31.66)=2.936, RM ANOVA) and *post-hoc* showed a significant TLS effect at (in min) 4 (*P*=0.0199), 6 (*P*=0.0255), 8 (*P*=0.0245), and 12 (*P*=0.0255) (**Fig.6h**).

When the OTR antagonist, OTA, was applied before TLS in response to 80 pA current injection, there was no TLS effect on firing frequency (*P*=0.2863, F (3.286, 23.00)=1.339, n=3 Type I and n=5 Type III neurons, RM ANOVA, **Fig.6h’**). As TLS evoked the most significant increase in firing at 4 min following stimulation, we also compared firing frequency with and without OTA at baseline vs. 4 min in response to 80 pA current for all neurons recorded. There was a significant TLS effect (*P*=0.0055, F(1, 17)=10.09, n=19, 2-way RM ANOVA), with no treatment effect (*P*=0.1705, F(1, 17)=2.048), and no interaction (*P*=0.3088, F(1, 17)=1.101). *Post-hoc* showed that TLS increased firing of BNST_DL_ neurons (*P*=0.0093) but had no effect when OTRs were blocked (*P*=0.3277, **Fig.6h’’**).

#### 3.9.2. AVP from the SCN fibers excites Type III BNST_DL_ neurons via OTR

In Type I BNST_DL_ neurons from rats with ChR2-eYFP expression in the SCN, there was a current effect on spike frequency (*P*=0.0001, F(1.338, 10.70)=28.60, n=9, two-way RM ANOVA), but no TLS effect (*P*=0.3718, F(1.888, 15.10)=1.045), nor interaction (*P*=0.2841, F(3.111, 24.89)=1.340). Similarly, when firing in response to 80 pA was analyzed separately, there was no TLS effect (*P*=0.5633, F(2.918, 23.34)=0.6903, n=9).

However, in Type III BNST_DL_ neurons, there was a trend in TLS effect on spike frequency (*P*=0.0586, F(1.383, 8.297)=4.443, 2-way RM ANOVA, n=7), and a significant current effect (*P*=0.0088, F(1.051, 6.306)=13.70), with no interaction (*P*=0.1178, F(2.840, 17.04)=2.286. *Post-hoc* analysis showed a significant TLS effect in response to 80-pA injection at (in min) 12 (vs. baseline *P*=0.0364), 14 (*P*=0.0154), 16 (*P*=0.0289), and 20 (*P*=0.0243). We then analyzed the response to 80-pA current separately and found a significant TLS effect (*P*=0.0251, F(1.739, 10.43)=5.611, n=7, RM ANOVA), with multiple comparisons showing significant TLS effects as above (**Fig.7d**).

In the presence of the OTR antagonist, OTA, TLS had no effect on firing of Type III neurons in response to 80-pA current injection (*P*=0.4261, F(1.945, 5.836)=0.9831, RM ANOVA, n=4, **Fig.7d’**). When Type III neurons’ firing following TLS (at 80pA) was analyzed with and without OTA, 2-way ANOVA showed a significant interaction between the TLS and OTA treatment (*P*=0.0059, F(1, 9)=12.84, n=11, 2-way RM ANOVA), with no TLS (*P*=0.7535, F(1, 9)=0.1048) or treatment effect (*P*=0.4243, F(1, 9)=0.7004). *Post-hoc* analysis showed that TLS increased firing of Type III neurons (*P*=0.0202), but this effect was not observed in the presence of OTA (*P*=0.1377, **Fig.7d’’**). We chose this time point as TLS evoked the most significant increase in firing at 14 min following stimulation.

#### 3.9.3. AVP from the PVN fibers excites a subset of III BNST_DL_ neurons via OTR

In Type I and III BNST_DL_ neurons from rats with ChR2-eYFP expression in the PVN, there was a current effect on spike frequency (*P*=0.0001, F(1.338, 10.70)=28.60, n=9, two-way RM ANOVA), no TLS effect (*P*=0.3718, F(1.888, 15.10)=1.045), and no interaction (*P*=0.2841, F(3.111, 24.89)=1.340). When spiking in response to 80 pA was analyzed separately, there was no TLS effect (*P*=0.3404, F(1.375, 12.38)=1.092, n=11, **Fig.7h’**). Similarly, there was no TLS effect in a presence of OTA (*P*=0.2920, F(1.668, 6.671)=1.457, n=5, **Fig.7h’’**). No treatment effect (*P*=0.4868, F(1, 14)=0.5102), TLS effect (F(1, 14)=1.145, *P*=0.3028), nor interaction (*P*=0.5684, F(1, 14)=0.3412, n=16) emerged when Type I and III neurons’ firing following TLS (at 80 pA) was analyzed with and without OTA (**Fig.7h’’’)**. However, two out of seven Type I BNST_DL_ neurons, both from rats with a very high bilateral PVN ChR2-eYFP expression, showed significant spiking increase to 283 and 144% of baseline at 6 min post TLS, whereas no Type I neurons showed increase in firing frequency when OTA was applied before TLS (**Fig.7h’’’**).

## 4. Discussion

We provide evidence that AVP-OT crosstalk via OTR-expressing BNST_DL_ neurons modulates defensive and exploratory behavior. Exogenous AVP directly and robustly excited Type I and Type III neurons in the BNST_DL_, a region critical for fear processing and vigilant threat monitoring ^25–28^, and this excitatory effect required OTR transmission. Specifically, OTR, but not V1aR or V1bR antagonists, blocked the excitatory effects of AVP on intrinsic membrane properties and firing frequency of BNST_DL_ neurons. In a subset of Type III BNST_DL_ neurons, V1aR activation by a selective V1aR agonist FE201874, which also blocks OTR ^47^, moderately increased firing without affecting intrinsic membrane properties. Excitatory effects of OTR have been documented in the cortex, hippocampus, and BNST ^40,63–65^, and both OTR and V1aR show excitatory effects in the central amygdala (CeA), though on different neuronal populations ^66–68^. AVP and OT differ by only two amino acids, and the cross-talk between these neuropeptides and their respective receptors has been well documented ^69–71^. OTR and V1aR, both centrally located G protein-coupled receptors (GPCRs) for OT and AVP ^72^, share 85% structural homology ^73,74^, while V1bR shares 40% identity with V1aR ^10^. Although OT and AVP have a similar affinity for OTR, OT has a higher affinity for OTR than for V1aR or V1bR ^75,76^.

The greater excitatory response of Type III neurons to AVP, which acts via OTR and V1aR, than to the endogenous OTR agonist, OT, or the highly selective and potent OTR agonist, TGOT, which acts via OTR alone ^77,78^, may be explained by the combined action of these receptors as well as the differences in the affinities of the neurohypophysial hormones toward the receptors. While AVP activates both OTR and V1aR signaling in Type III BNST_DL_ neurons to robustly increase excitability, in Type I neurons, V1aR appears to provide tonic inhibition to a subset of neurons, despite the excitatory effect of OTR. This inhibition was unmasked by the excitatory effect of V1aR antagonist (SR49059), and the inhibitory effects of central V1aR transmission have been reported before ^79^. However, OTR remains the key player in regulating BNST_DL_ neuron excitability and BNST_DL_- dependent behaviors, which are directly influenced by AVP-OT crosstalk.

In many brain regions, the impact of OT and AVP crosstalk on physiology and behavior is less pertinent because OTR and V1R are frequently located in anatomically separate neuronal circuits. In these cases, the effects of OT and AVP depend on the brain site-specific expression of their receptors ^80,81^. However, the BNST_DL_ expresses both OTR ^82,83^ and V1aR ^84^ and here, we show cell type-specific functional expression of these receptors in Type I and III neurons. This finding aligns with our recent study showing postsynaptic expression of OTR in these neuron types ^40^. We further confirmed the excitatory effect of AVP on fluorescent OTR-BNST_DL_ neurons, with the great majority of OTR neurons characterized as Type III. Furthermore, previous single-cell PCR studies have shown that Type III BNST_DL_ neurons express OTR, CRF, and Striatal Enriched Protein Tyrosine Phosphatase (STEP) mRNA ^33^. Using CRF-Cre transgenic rats, we showed that recorded CRF-mCherry-BNST_DL_ neurons, which exhibit electrophysiological properties of Type III neurons, are directly excited by both AVP and TGOT. While the co-expression of CRF and OTR in BNST_DL_ neurons was first described in 2011 ^33^, this study provides the first functional evidence of OTR activation directly exciting CRF-BNST_DL_ neurons. Consistent with previous *in situ* hybridization and OTR binding studies ^18,82,85^, identifying the BNST as one of the highest OTR-expressing regions in the rodent brain, we found numerous OTR-expressing neurons throughout the BNST of OTR-Cre rats. Recent single-cell RNAseq analysis in mice revealed that OTRs are expressed in diverse populations of GABA-ergic neurons in the BNST^86^. Here, we found that OTR-BNST_DL_ neurons co-expressed either PKCδ or STEP, two enzymes present on mutually exclusive populations of neurons in the BNST_DL_ ^58^. STEP is a unique marker of CRF neurons in the oval nucleus of the BNST_DL_ and is exclusively expressed by Type III BNST_DL_ neurons on a single-cell level ^35^. While our study primarily focused on the BNST_DL_, different types of OTR neurons might be present in the posterior BNST. Our results contrast with the functional expression and roles of OTR and V1aR in the CeA, where these receptors are located on mutually exclusive neuronal populations in the lateral and medial CeA, respectively, where they play opposite roles in inhibiting and exciting medial CeA output neurons and fear behaviors ^16,53,63^. Thus, Type III BNST_DL_ neurons represent a unique cell population in the extended amygdala in which OTR and V1aR signaling converge to increase excitability in response to AVP.

OTR signaling in the extended amygdala plays an intricate role in fear and vigilance. OTR activation in the CeA reduced contextual fear ^53,87,88^, whereas OTR in the basolateral nucleus of the amygdala (BLA) strengthened fear discrimination toward discrete cues predicting danger ^89^. Our previous behavioral studies showed that blocking OTR transmission in the BNST_DL_ reduced the acquisition and recall of learned cued fear measured in the FPS ^27^. Interestingly, blocking OTR in the BNST_DL_ also reduced the ratio between cued and non-cued fear in the same rats, suggesting that BNST_DL_ OTR transmission strengthens fear responses to discrete (cued) threats while reducing threat-induced anxious arousal ^27,39^. Systemic or intracerebroventricular OT similarly reduces non-cued fear in the FPS ^90–92^. In this study, chemogenetic silencing of OTR-BNST_DL_ neurons prolonged the expression of non-cued fear measured in FPS. The non-cued fear reflects potentiation of the startle response that occurs between cue presentations during the cued fear recall, emerging only after the cue is being presented ^27,91^. Hence, this response is thought to reflect a cue-induced vigilant state or anxious arousal, which is independent from contextual fear ^93^. These findings underscore the unique role of OTR and OTR-BNST_DL_ neurons in biasing fear learning toward imminent and signaled threats while dampening threat-induced anxious arousal.

We also found that in the fear-conditioned rats, OTR-BNST_DL_ neuron activity increased open-arm exploration time in the EPM. This aligns with extensive literature showing anxiolytic-like effects of OTR activation in the EPM, especially in stressed animals, for review see ^7,8,94^. BNST_DL_ activity is generally thought to promote anxiety-like behavior by mediating hypervigilance and anticipatory anxiety in response to unpredictable or un-signaled threats ^26,95,96^. Although this appears in contrast with the role of OTR-BNST_DL_ neurons, our recent study demonstrated that OTR activation excites BNST_DL_ interneurons, thereby increasing inhibitory synaptic transmission that inhibits Type II BNST_DL_ output neurons. This results in a primarily inhibitory effect of OTR activation on BNST_DL_ output to the CeA ^83^, a projection which has been shown before to reduce open arms exploration in the EPM ^97^.

AVP and OT influence fear- and anxiety-like behaviors, and are often labeled as ‘anxiogenic’ and ‘anxiolytic’ peptides, respectively. The anxiogenic label for AVP originates from early studies on AVP-deficient Brattleboro rats, which showed reduced vigilance in open spaces and reduced contextual fear memory ^98–100^. Consistent with this, systemic V1aR activation reduces exploration of open arms in the EPM, while V1aR antagonism or knockout increases exploration ^19,101–103^. However, V1aR activation in the lateral septum and posterior BNST can increase EPM exploration and reduce defensive behaviors ^31,104^, suggesting a more complex role of V1aR in modulating fear behavior. Notably, the commonly used V1aR-selective antagonist, d(CH_2_)_5_[Tyr(Me)^2^]AVP (Manning compound)^19,20^, though not binding V1bR or V2R, acts as a potent OTR antagonist ^49^. Given the abundant expression of OTR and V1R in the hypothalamus and extended amygdala, the behavioral effects of AVP via OTR may be underestimated. Our finding that AVP increased BNST_DL_ neuronal excitability, requiring OTR signaling (with V1aR showing a moderate excitatory effect), suggests that OTR and V1aR, modulated by specific hypothalamic inputs, work in tandem rather than in opposition to impact BNST neuron activity and behaviors. However, since we used blockers of synaptic transmission in the current study, we cannot exclude the possibility that V1aR, similarly to OTR ^34,40^, has a distinct effect on synaptic activity in the BNST_DL_.

This study has other limitations. Firstly, female rats were not included. Although we aim to explore potential sex differences (or the lack thereof) in the effects of the AVP and OT systems on BNST excitability and behavior, including both sexes for a detailed analysis was beyond the scope of this study. Combining males and females in the same experimental groups could obscure the significant role the BNST_DL_ OTR and V1aR ^29,105^ play in mediating sex differences and similarities in stress-related behaviors, including non-cued fear, which also differs between male and female rats^93^. Additionally, we cannot rule out the possibility that local AVP neurons from the posterior BNST, which exhibit strong sex-dependent expression and function (with high levels in male rats) ^106^, may also project to the BNST_DL_ ^23^.

Previous studies have shown that the rat BNST_DL_ receives OT inputs from the PVN ^33,39,53^, which we confirmed in the current study using optogenetic approach. We also showed that the BNST_DL_ receives AVP inputs from the SON, SCN, and the PVN and that optogenetic high-frequency stimulation of AVP terminals in the BNST_DL_ increased Type I and III BNST_DL_ neurons’ excitability via OTR. Cell-selective genetic manipulations have facilitated our understanding of the contributions of specific populations of hypothalamic and extra-hypothalamic AVP neurons to defensive behaviors. For example, eliminating AVP neurons in the PVN or SCN using caspase-mediated methods in AVP-Cre mice reduced open-arm exploration in the EPM ^107^, whereas removing AVP cells in the posterior BNST did not influence open-arm exploration. Overall, these findings suggest that hypothalamic AVP neurons diminish vigilance and promote open-space exploration in the EPM. Although the specific role of AVP SON neurons in defensive behavior is not yet known, it is noteworthy that the activity of AVP neurons in the SCN ^107^ and the activity of OTR neurons in the BNST_DL_ both increase rodents’ open-arm exploration time in the EPM. Thus, in periods when AVP neuron activity in the SCN is high, such as during preparation for the rest phase ^1^, the system may be primed for AVP release in the BNST_DL_ in response to physiological stimuli, which would activate OTR neurons in the BNST_DL_ and reduce defensive behaviors. Our study reveals cell type and receptor-specific modulation of BNST_DL_ neuron activity via unique hypothalamic projections containing neurohypophysial hormone AVP. We showed that although both OT and AVP have excitatory effects on Type III OTR-expressing neurons of the BNST_DL_, they differed in the receptors that contributed to the response (OTR alone vs. OTR and V1aR) and hence, the magnitude of the excitatory response. These results suggest that changes in internal states favoring the release of OT vs. AVP (e.g. lactation vs. thirst/salt intake) will have a scaled excitatory impact on Type III OTR-expressing neurons of the BNST_DL_ and will fine tune their input-output responses, thus providing them with greater flexibility in response to specific neurohypophysial inputs.

## 5. Acknowledgements

This work was supported by grant MH113007 from National Institute of Mental Health (NIMH) to JD and by the Synergy European Research Council (ERC) grant “OxytocINspace” 101071777, SFB Consortium 1158-3, and German-Israeli Project cooperation (DIP) GR3619-1 to VG.

We thank Dr. Robert Messing, University of Texas Austin for providing us with the CRF-Cre transgenic rats breeders.

We would like to thank Dr. Sarah Daniel of In Scripto, LLC, for edits and comments on this manuscript.

## 7. Authors contributions

Conceptualization: JD; Methodology: WF, VOP, FB, SLO, QK, VG, JD; Formal analyses: FB, WF, VOP, RC, JD; Investigation: WF, VOP, FB, SLO, RC, LMM, QR, JD; Resources: QK, VG, JD; Writing - original draft: WF, VOP, JD; Writing – review and editing: WF, VOP, FB, RC, VG, JD; Visualization: JD; Supervision: JD; Project administration: JD; Funding acquisition; VG, JD

## Supplementary Materials

**Supplementary Table 1.** Pharmacological compounds used to manipulate OTR, V1aR, and V1bR.

| Compound | Alternative name, Abbreviation | Company Catalog # | Concentration | Receptor activity profile |
| --- | --- | --- | --- | --- |
| Arginine-vasopressin | AVP, Arg<sup>8</sup>-vasopressin | Bachem Americas, #4012215 | 0.2 µM | V1R, V2, and OTR agonist<sup>66</sup> |
| Oxytocin | OT | Bachem Americas, #4016373 | 0.2 µM | OTR and V1R agonist<sup>108</sup> |
| (Thr<sup>4</sup>,Gly<sup>7</sup>)-Oxytocin H-Cys-Tyr-Asn-Cys-Gly-Leu-Gly-NH<sub>2</sub> | TGOT | Bachem Americas, 4013837 | 0.4 µM | Selective OTR agonist<sup>65</sup> |
| FE 201874 |  | Ferring Pharmaceuticals | 0.2-0.4 µM | Selective V1aR agonist, OTR antagonist<sup>109</sup> |
| d(Cha4)-AVP |  | GLPBIO, #GC1659 | 1 µM | Selective V1bR agonist<sup>110</sup> |
| (d(CH<sub>2</sub>)<sub>5</sub><sup>1</sup>,Tyr(Me)<sup>2</sup>, Arg<sup>8</sup>)-vasopressin | Manning compound | Tocris, Bio-Techne Corporation, #3377 | 1 µM | V1aR/OTR receptor antagonist<sup>111</sup> |
| SR 49059 | Relcovaptan | Tocris, Bio-Techne Corporation, #2310 | 5 µM | Selective V1aR antagonist<sup>112</sup> |
| d(CH<sub>2</sub>)<sub>5</sub>(1), D-Tyr(2), Thr(4), Orn(8), des-Gly-NH<sub>2</sub>(9)]-Vasotocin | OTA | Chemical Repository, #V-905 NIMH | 0.4 µM | Selective OTR antagonist<sup>110</sup> |
| SSR 149415 | Nelivaptan | Tocris, Bio-Techne Corporation, #6195 | 1 µM | Selective V1bR antagonist<sup>113</sup> |

### Supplementary results

#### 1. The effect of AVP on intrinsic membrane properties of Type I BNST_DL_ neurons

We first determined the effect of AVP on membrane properties of Type I-III BNST_DL_ neurons (see Table 1). In Type I neurons, AVP induced a significant depolarization of the RMP (mV) from −59.1±1.77 to −53.87±1.25 (*P*=0.0101, F(2, 15)=6.340), n=9, mixed-effect model). This change was associated with a significant increase in Rin (MΩ; at −100 pA) from 139.54±11.74 to 165.42±16.62 (*P*=0.0187, F(1.774, 13.31)=5.695, n=9). AVP induced a significant reduction in Rh (pA) from 27.08±3.24 to 15.42±2.01 (*P*=0.0072, F(2, 13)=7.374, n=8). The first-spike Th (mV) was not significantly changed from −35.83±0.79 to −35.80±0.65 after AVP (*P*=0.4681, F(1.300, 9.750)=0.6786, n=9), but the first-spike Lat (ms) was significantly reduced from 221.50±56.41 to 137.50±33.78 (*P*=0.0058, F(2, 15)=7.393, n=9, mixed-effect analysis).

In the presence of the OTR antagonist, OTA, AVP did not significantly affect the Rh (pA) at 36.58±4.16 and 30.17±4.73 (*P*=0.1840, F(1.584, 14.26)=1.938, n=10), or first-spike Th (mV) at −35.38± 0.71 and −35.72±0.88 (*P*=0.6484, F(1.720, 15.48)=0.3976, n=10). The first-spike Lat (ms), however, was significantly reduced from 188.20±33.61 to 134.30±32.14 (*P*=0.0293, F(1.967, 17.70)=4.366, n=9, RM one-way ANOVA), but multiple comparisons only showed a trend in differences between the groups (*P*=0.0599, pre vs. OTA+AVP). There was also a treatment effect on the RMP (in mV) from - 63.17±1.48 to −57.04±2.15 (*P*=0.0244, F(1.088, 9.788)=6.851, n=10), but multiple comparisons showed that this effect was driven by OTA alone (*P*=0.0497, pre vs. OTA). Lastly, Rin (MΩ) (at −100 pA) was increased from 191.60±15.27 to 202.60±14.01 (*P*=0.0162, F(1.719, 15.47)=5.789, n=10), but multiple comparisons showed only a trend for the AVP effect (*P*=0.0575, pre vs. OTA+AVP).

In contrast, AVP application in a presence of V1aR antagonist, SR49059, induced a significant depolarization of RMP (mV) from −64.22±1.93 to −62.47±2.12 (*P*=0.0240, F(1.571, 12.57)=5.548, n=9, RM ANOVA), and the Rh (pA) from 45.37±7.00 to 38.61±6.76 (*P*=0.0120, F(1.806, 14.45)=6.361). However, it did not affect Rin (MΩ) from 180.1±15.50 to 202.1±16.27 (−100 pA, *P*=0.0930, F(1.216, 9.730)=3.353), the first-spike Lat (ms) from 161.1±18.26 to 143.4±21.08 (*P*=0.4393, F(1.273, 10.19)=0.7476), or Th (mV) from −36.23±1.51 to −34.88±1.72 (*P*=0.1401, F(1.260, 10.08)=2.523).

AVP application in the presence of the V1bR antagonist Nelivaptan significantly reduced the RMP (mV) from −63.13±2.28 to −56.76±2.73 (*P*=0.0010, F(1.352, 12.17)=15.50, n=10, RM one-way ANOVA) and the Rh (pA) from 39.50±5.70 to 31.00±5.36 (*P*=0.0093, F(1.387, 12.48)=8.143, n=10). It also increased the Rin (MΩ) (at −250 pA) from 189.5±16.30 to 214.6±17.92 (*P*=0.0480, F(1.421, 12.79)=4.299, n=10). While there was no effect on first-spike Th (mV) at −30.28±1.69 and −28.59±2.22 (*P*=0.1335, F(1.292, 11.63)=2.541, n=10), the first-spike Lat (ms) was reduced from 187.2±28.40 to 113.6±18.15 (*P*=0.0090, F(1.707, 15.37)=6.963, n=10) after Nelivaptan and AVP application.

#### 2. The effect of AVP on membrane properties of Type II BNST_DL_ neurons

AVP did not induce a significant depolarization of the RMP (mV) in Type II BNST_DL_ neurons from - 53.12±1.46 to −51.29±1.69 (*P*=0.1815, F(1.155, 6.927)=2.219, n=8, mixed-effect analysis). Rin (MΩ) (at −100 pA) remained unchanged at 135.9±15.83 and 151.3±21.57 (*P*=0.0699, F(2, 12)=3.348, n=8), as did Rh (pA) at 25.10±5.39 and 25.62±8.73 (*P*=0.7867, F(1.016, 6.095)=0.08300, n=8). Similarly, neither the first-spike Th (mV) at −36.82±0.82 and −34.65±1.53 (*P*=0.3336, F(1.818, 10.91)=1.199, n=8), nor the first-spike Lat (ms) at 97.3±7. 3 and 75.27±8.7 (*P*=0.3397, F(1.403, 8.415)=1.146, n=8) were changed after AVP.

#### 3. The effect of AVP on membrane properties of Type III BNST_DL_ neurons

In Type III neurons, AVP induced a significant depolarization of RMP (mV) from −68.34±1.36 to - 65.±1.93 (*P*=0.0143, F(2, 15)=5.717, n=10, mixed-effect analysis), associated with a significant increase in Rin (MΩ; at −100 pA) from 108.8±10.29 to 128.9±13.83 (*P*=0.0011, F (1.602, 5.608)=30.90, n=5). Consequently, the Rh (pA) was significantly reduced from 75.15±9.45 to 63.5±9.65 (*P*=0.0068, F(1.114, 8.356)=12.18, n=10). The first-spike Th (mV) was not changed, at - 33.12±1.89 and −33.46±1.92 before and after AVP, respectively (*P*=0.5495, F(1.187, 8.902)=0.4559, n=10), but AVP induced a significant reduction in the first-spike Lat (ms) from 419.08±64.09 to 257.5±52.13 (*P*=0.0014, F(1.869,13.08)=11.79, n=10).

Next, we applied a series of OTR/V1R antagonists to determine whether the effects of AVP were mediated via V1aR, V1bR, or OTR. In the presence of the OTR/V1aR antagonist ((d(CH_2_)_5_^1^,Tyr(Me)^2^,Arg^8^)-Vasopressin), AVP did not affect the RMP (mV) at −69.79±1.21 and - 67.97±1.17 (*P*=0.0936, F(1.211, 7.264)=3.616, n=7, RM one-way ANOVA), Rin (MΩ) (at −100 pA) at 94.74±16.92 and 97.37±17.56 (*P*=0.1761, F(1.904, 11.42)=2.037, n=7), Rh (pA) at 88.33±8.97 and 83.33±4.42 (*P*=0.4213, F(1.335, 8.007)=0.8376, n=7), first-spike Th (mV) at −33.41±0.95 and - 34.69±0.79 (*P*=0.1517, F(1.344, 8.063)=2.478, n=7), or first-spike Lat (ms) at 203.2±29.08 and 176.2±32.8’1 (*P*=0.5843, F(1.174, 7.041)=0.3909, n=7) before and after AVP application, respectively.

Further, in the presence of OTA, AVP did not significantly affect RMP (mV) at −67.37±1.72 and - 66.59±1.9 (*P*=0.4675, F(1.039, 6.235)=0.6159, n=7, RM one-way ANOVA), Rin (MΩ, at −100 pA) at 110.99±8.5 and 132.79±15 (*P*=0.0758, F(1.042, 4.169)=5.504, n=5), Rh (pA) at 77.02±10.75 and 75.71±12.74 (*P*=0.8495, F(1.197, 7.185)=0.06355, n=7), first-spike Th (mV) at −38.86±1.96 and - 37.58±2.46 (*P*=0.2667, F(1.926, 11.56)=1.482, n=7, RM one-way ANOVA), or first-spike Lat (ms) at 327.56±60.45 and 300.3±60.50 (*P*=0.5761, F(2, 10)=0.5830, n=6) before and after AVP application, respectively.

However, in the presence of the V1aR antagonist, SR49059, AVP induced a significant depolarization of RMP (mV) from −66.95±1.88 to −63.94±2.38 (*P*=0.0052, F(1.341, 9.386)=11.51, n=8, RM one-way ANOVA) and a significant increase in Rin (MΩ) (at −100 pA) from 159±13.93 to 181.5±14.85 (*P*=0.0179, F(1.764, 12.35)=5.944, n=8), as well as a significant decrease in Rh (pA) from 68.96±7.48 to 56.67±7.13 (*P*=0.0193, F(1.249, 8.745)=7.536, n=8). The first-spike Lat (ms) was also significantly decreased when AVP was applied in the presence of the V1aR blocker, from 404.0±72.91 to 261.2±65.96 (*P*=0.0121, F(1.546, 10.83)=7.551, n=8). Consistent with the effects of AVP alone, AVP in the presence of SR49059 did not change the first-spike Th (mV), at −34.09±1.33 and −33.25±2.33 (*P*=0.2014, F(1.844, 12.91)=1.825, n=8) before and after AVP, respectively.

Lastly, in the presence of the V1bR antagonist, Nelivaptan, AVP induced a significant depolarization of RMP (mV) from −74.24±1.46 to −69.45±1.74 (*P*=0.0014, F(1.331, 10.65)=15.67, n=9, RM one-way ANOVA) and a significant increase in Rin (MΩ) (at −250 pA) from 124.9±9.76 to 186.6±24.80 (*P*=0.0114, F(1.196, 9.565)=9.060, n=9), as well as a significant decrease in Rh (pA) from 94.44±5.86 to 84.26±6.50 (*P*=0.0272, F(1.507, 12.06)=5.441, n=9). The first-spike Lat (ms) was significantly reduced when AVP was applied in the presence of the V1bR antagonist, Nelivaptan, from 377.2±64.25 to 248.4±37.86 (*P*=0.0420, F(1.539, 12.31)=4.491, n=9). Consistently with the effects of AVP alone, the first-spike Th (mV) was not changed, at −29.90±1.76 and −24.24±3.12 (*P*=0.0610, F(1.077, 8.613)=4.558, n=9) before and after AVP application, respectively.

These results suggest that AVP excites Type III BNST_DL_ neurons via a direct postsynaptic mechanism that involves OTR but not V1aR or V1bR activation.

#### 4. The effect of OT on membrane properties of Type III BNST_DL_ neurons

OT did not induce a significant depolarization of the RMP (mV) in Type III BNST_DL_ neurons, at - 68.54±1.03 and −67.79±0.87 (*P*=0.2875, F(2, 13)=1.374, n=8, mixed-effect analysis). Rin (MΩ) (at - 100 pA) significantly changed from 137.15±23.61 to 142.41±26.16 (*P*=0.0251, F(0.8062, 3.628)=14.14, n=6), as did Rh (pA), at 83.02±4.81 and 69.33±4.74 (*P*=0.0007, F(2, 13)=13.22, n=8). Similarly to the AVP effect, the first-spike Th (mV) was not changed by OT at −29.27±1.86 and - 29.49±1.98 (*P*=0.1480, F(0.04181, 0.2717)=2.479, n=8) but the first-spike Lat (ms) significantly changed with OT application from 497.8± 63.95 to 281.05±43.37 (*P*=0.0027, F(1.827, 10.96)=11.13, n=8).

#### 5. The effect of OTR agonist, TGOT, on membrane properties of Type III BNST_DL_ neurons

The results above imply that AVP affects the membrane properties of Type III BNST_DL_ neurons via OTR. Thus, we next applied a potent OTR agonist, TGOT. TGOT induced a significant depolarization of the RMP (mV) from −71.86±1.00 to −70.00±1.22 (*P*=0.0469, F(1.168, 8.177)=5.241, n=9, mixed-effect analysis) and a significant increase in Rin (MΩ) (at −100 pA) from 85.79±6.63 to 91.24±7.02 (*P*=0.0338, F(1.076, 4.841)=8.456, n=7). TGOT significantly decreased Rh (pA) from 93.61±9.88 to 82.69±10.97 (*P*=0.0174, F(1.776, 11.54)=6.129, n=9) and decreased the first-spike Lat (ms) from 594.03±76.46 to 355.243±68.69 (*P*=0.0373, F(0.9019, 5.412)=7.726, n=9). Similarly to the effect of AVP and OT, TGOT did not significantly change the first-spike Th (mV), at −30.84±0.98 and - 30.79±1.13 (*P*=0.4439, F (1.120, 7.838)=0.6998, n=9) before and after TGOT, respectively. These results confirm the primary contribution of OTR to the effects of AVP on membrane properties of Type III BNST_DL_ neurons.

#### 6. The effect of V1aR agonist, FE 201874, on membrane properties of Type III BNST_DL_ neurons

In contrast to TGOT, application of the selective rat V1aR agonist, FE 201874, did not change the RMP (mV) at −70.72±0.9 and −69.44±1.25 (*P*=0.1068, F(1.242, 9.312)=3.093, n=10, mixed-effect analysis), Rin (MΩ) (at −250 pA) at 118.21±1.25 and 117.57±9.69 (*P*=0.70748, F(1.219, 6.094)=0.2128, n=7, mixed-effect analysis), Rh (pA) at 77.83±9.55 and 73.66±10.5 (*P*=0.2256, F(1.209, 9.066)=1.728, n=10), first-spike Lat (ms) at 432.8±65.72 and 345.3±57.68 (*P*=0.0610, F(1.710, 11.12)=3.780, n=9), or first-spike Th (mV) at −30.45±1.22 and −31.93±1.1 (*P*=0.1312, F(1.550, 11.63)=2.515, n=10) before and after FE 201874 application, respectively.

#### 7. The effect of V1bR agonist, d[Cha4]-AVP, on membrane properties of Type III BNST_DL_ neurons

V1bR agonist, d[Cha4]-AVP, did not change the RMP (mV) in Type III BNST_DL_ neurons, with a RMP of −70.42±0.84 and −69.81±1.15 pre and post V1bR agonist application, respectively (*P*=0.2988, F(1.987, 12.91)=1.327, n=8, mixed-effect analysis). Rin (MΩ) (at −250 pA) remained unchanged at 105.18±7.85 and 110.98±10.05 (*P*=0.1532, F(0.9314, 6.054)=2.663, n=8), as did Rh (pA), at 86.25±5.96 and 81.46±6.2 (*P*=0.3195, F(1.160, 7.540)=1.193, n=8). Similarly, the first-spike Th (mV) was not significantly changed at −33.38± 1.67 and −34.00±1.79 by the V1bR agonist (*P*=0.1666, F(0.9257, 6.017)=2.467, n=8). However, the first-spike Lat (ms) was significantly changed from 475.1±59.48 to 426.6±60.20 before and after V1bR agonist (*P*=0.0185, F(1.063, 5.848)=10.24, n=8).

**Supplementary figure 1.**
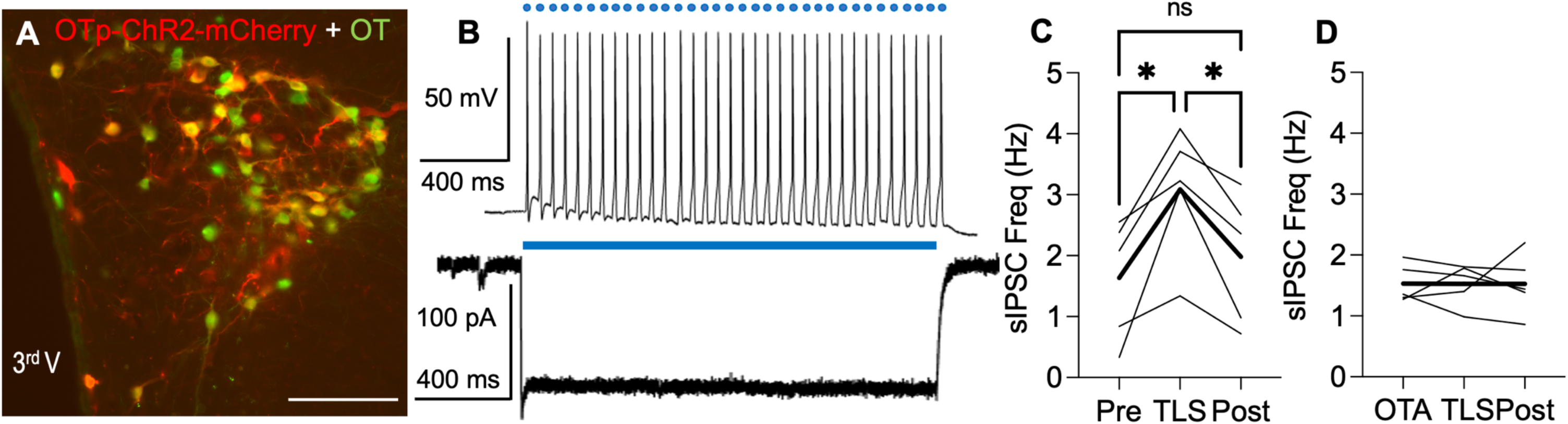
TLS evokes OT release and increases inhibitory synaptic transmission in Type II BNST_DL_ neurons in an OTR-dependent manner. **A**: High somatodendritic expression of ChR2-mCherry (red) is observed in hypothalamic OT neurons (mouse anti-OT MAB5296 antibody, green) 3 weeks after *AAV-OTp-ChR2-mCherry* was injected to the PVN in male rats (20x, scale bar 100 μm). **B**: Action potentials recorded from PVN neurons expressing ChR2-mCherry, evoked by 10 ms repeated (30 Hz, 1 sec) blue light stimulation (upper panel) and current induced by 1 sec continuous blue light (bottom panel). **C**: In the BNST_DL_ from rats injected as above, TLS (30 Hz train, single pulse duration 10 ms for 20 sec) increased frequency of spontaneous inhibitory postsynaptic currents (sIPSCs) in Type II BNST_DL_ neurons (F(1.436, 7.182)=16.86, *P*=0.0028, one-way ANOVA, n=5) but did not affect sIPSCs amplitude (F(1.851, 7.404)=2.471, *P*=0.1520, not shown). This effect mimics a well-established cellular effect of exogenous OT in the BNST_DL_, which was shown to increase frequency of sIPSCs specifically in Type II BNST_DL_ neurons ^40^. **D**: Notably, TLS did not affect sIPSCs frequency in the presence of selective OTR antagonist, OTA (F(1.565, 7.823)=0.0003, *P*=0.9985, n=5). Fine lines represent individual Type II neurons responses, black thick lines - average responses, * *P*<0.05, 3^rd^ V - Third ventricle.

**Supplementary figure 2.**
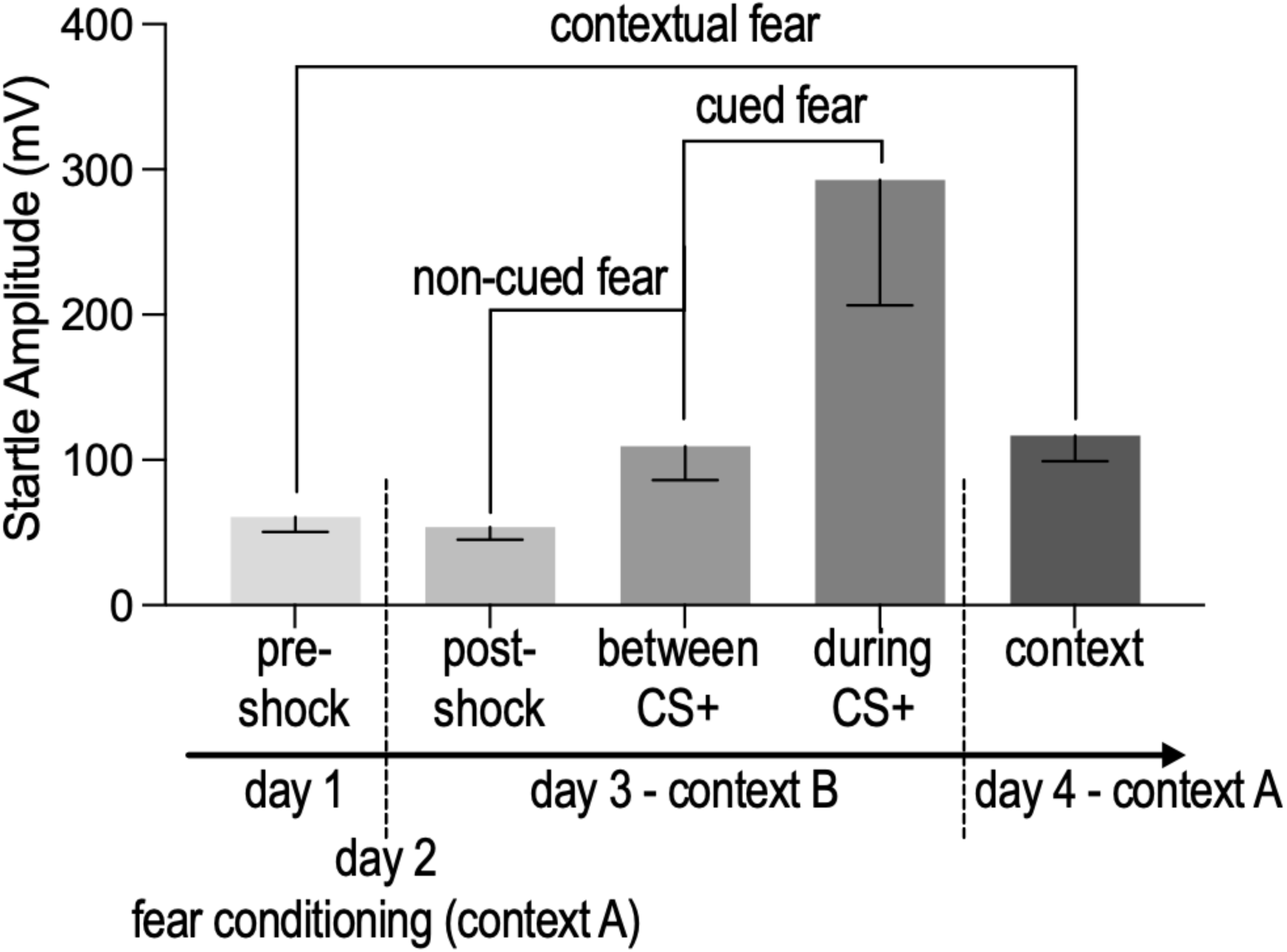
Fear-potentiated startle (FPS) components. Rats are tested for baseline acoustic startle response ASR, pre-shock, day 1) before fear conditioning on day 2 (context A). On day 3 fear recall is tested in context B: <u>Cued fear</u> is calculated as a percent change score of ASR amplitude from ‘between conditioned stimulus (CS+)’ to ASR ‘during CS+’. <u>Non-cued fear</u> is calculated as a percent change score from post-shock to ‘between CS+’. <u>Note</u>, during the fear recall test, post-shock ASR is measured before the first CS+ presentation. On day 4 rats are tested for <u>contextual fear</u> recall in context A with no CS presentations, calculated as percent change score from pre-shock on day 1 to ASR in context A on day 4.

## References

1. Kalsbeek, A., Fliers, E., Hofman, M.A., Swaab, D.F., and Buijs, R.M. (2010). Vasopressin and the output of the hypothalamic biological clock. J Neuroendocrinol 22, 362–372. 10.1111/j.1365-2826.2010.01956.x.

2. Brown, C.H., Ludwig, M., Tasker, J.G., and Stern, J.E. (2020). Somato-dendritic vasopressin and oxytocin secretion in endocrine and autonomic regulation. J Neuroendocrinol 32, e12856. 10.1111/jne.12856.

3. Serradeil-Le Gal, C., Wagnon, J., Tonnerre, B., Roux, R., Garcia, G., Griebel, G., and Aulombard, A. (2005). An overview of SSR149415, a selective nonpeptide vasopressin V(1b) receptor antagonist for the treatment of stress-related disorders. CNS Drug Rev 11, 53–68. 10.1111/j.1527-3458.2005.tb00035.x.

4. Caldwell, H.K., Lee, H.-J., Macbeth, A.H., and Young, W.S. (2008). Vasopressin: behavioral roles of an “original” neuropeptide. Prog Neurobiol 84, 1–24. 10.1016/j.pneurobio.2007.10.007.

5. Veenema, A.H. (2012). Toward understanding how early-life social experiences alter oxytocin- and vasopressin-regulated social behaviors. Horm Behav 61, 304–312. 10.1016/j.yhbeh.2011.12.002.

6. Jurek, B., and Neumann, I.D. (2018). The Oxytocin Receptor: From Intracellular Signaling to Behavior. Physiological Reviews 98, 1805–1908. 10.1152/physrev.00031.2017.

7. Janeček, M., and Dabrowska, J. (2019). Oxytocin facilitates adaptive fear and attenuates anxiety responses in animal models and human studies—potential interaction with the corticotropin-releasing factor (CRF) system in the bed nucleus of the stria terminalis (BNST). Cell Tissue Res 375, 143–172. 10.1007/s00441-018-2889-8.

8. Olivera-Pasilio, V., and Dabrowska, J. (2020). Oxytocin Promotes Accurate Fear Discrimination and Adaptive Defensive Behaviors. Front. Neurosci. 14, 583878. 10.3389/fnins.2020.583878.

9. Khan, S., Raghuram, V., Chen, L., Chou, C.-L., Yang, C.-R., Khundmiri, S.J., and Knepper, M.A. (2024). Vasopressin V2 receptor, tolvaptan, and ERK1/2 phosphorylation in the renal collecting duct. Am J Physiol Renal Physiol 326, F57–F68. 10.1152/ajprenal.00124.2023.

10. Lolait, S.J., O’Carroll, A.M., Mahan, L.C., Felder, C.C., Button, D.C., Young, W.S., Mezey, E., and Brownstein, M.J. (1995). Extrapituitary expression of the rat V1b vasopressin receptor gene. Proc Natl Acad Sci U S A 92, 6783–6787. 10.1073/pnas.92.15.6783.

11. Corbani, M., Marir, R., Trueba, M., Chafai, M., Vincent, A., Borie, A.M., Desarménien, M.G., Ueta, Y., Tomboly, C., Olma, A., et al. (2018). Neuroanatomical distribution and function of the vasopressin V1B receptor in the rat brain deciphered using specific fluorescent ligands. Gen Comp Endocrinol 258, 15–32. 10.1016/j.ygcen.2017.10.011.

12. Tribollet, E., Barberis, C., Jard, S., Dubois-Dauphin, M., and Dreifuss, J.J. (1988). Localization and pharmacological characterization of high affinity binding sites for vasopressin and oxytocin in the rat brain by light microscopic autoradiography. Brain Research 442, 105–118. 10.1016/0006-8993(88)91437-0.

13. Tribollet, E., Dubois-Dauphin, M., Dreifuss, J.J., Barberis, C., and Jard, S. (1992). Oxytocin receptors in the central nervous system. Distribution, development, and species differences. Ann. N. Y. Acad. Sci. 652, 29–38.

14. Johnson, A.E., Audigier, S., Rossi, F., Jard, S., Tribollet, E., and Barberis, C. (1993). Localization and characterization of vasopressin binding sites in the rat brain using an iodinated linear AVP antagonist. Brain Res 622, 9–16. 10.1016/0006-8993(93)90795-o.

15. Roozendaal, B., Schoorlemmer, G.H., Wiersma, A., Sluyter, S., Driscoll, P., Koolhaas, J.M., and Bohus, B. (1992). Opposite effects of central amygdaloid vasopressin and oxytocin on the regulation of conditioned stress responses in male rats. Ann N Y Acad Sci 652, 460–461. 10.1111/j.1749-6632.1992.tb34384.x.

16. Viviani, D., and Stoop, R. (2008). Opposite effects of oxytocin and vasopressin on the emotional expression of the fear response. Prog Brain Res 170, 207–218. 10.1016/S0079-6123(08)00418-4.

17. Veinante, P., and Freund-Mercier, M.J. (1997). Distribution of oxytocin- and vasopressin-binding sites in the rat extended amygdala: a histoautoradiographic study. J. Comp. Neurol. 383, 305– 325.

18. Smith, C.J.W., DiBenedictis, B.T., and Veenema, A.H. (2019). Comparing vasopressin and oxytocin fiber and receptor density patterns in the social behavior neural network: Implications for cross-system signaling. Frontiers in Neuroendocrinology 53, 100737. 10.1016/j.yfrne.2019.02.001.

19. Bayerl, D.S., Hönig, J.N., and Bosch, O.J. (2016). Vasopressin V1a, but not V1b, receptors within the PVN of lactating rats mediate maternal care and anxiety-related behaviour. Behav. Brain Res. 305, 18–22. 10.1016/j.bbr.2016.02.020.

20. Hernández-Pérez, O.R., Crespo-Ramírez, M., Cuza-Ferrer, Y., Anias-Calderón, J., Zhang, L., Roldan-Roldan, G., Aguilar-Roblero, R., Borroto-Escuela, D.O., Fuxe, K., and Perez de la Mora, M. (2018). Differential activation of arginine-vasopressin receptor subtypes in the amygdaloid modulation of anxiety in the rat by arginine-vasopressin. Psychopharmacology (Berl.) 235, 1015–1027. 10.1007/s00213-017-4817-0.

21. Rood, B.D., Stott, R.T., You, S., Smith, C.J.W., Woodbury, M.E., and De Vries, G.J. (2013). Site of origin of and sex differences in the vasopressin innervation of the mouse (Mus musculus) brain. J Comp Neurol 521, 2321–2358. 10.1002/cne.23288.

22. Brown, C.H., Ludwig, M., Tasker, J.G., and Stern, J.E. (2020). Somato-dendritic vasopressin and oxytocin secretion in endocrine and autonomic regulation. J Neuroendocrinol 32. 10.1111/jne.12856.

23. Rigney, N., de Vries, G.J., and Petrulis, A. (2023). Sex differences in afferents and efferents of vasopressin neurons of the bed nucleus of the stria terminalis and medial amygdala in mice. Horm Behav 154, 105407. 10.1016/j.yhbeh.2023.105407.

24. Mohr, E., Bahnsen, U., Kiessling, C., and Richter, D. (1988). Expression of the vasopressin and oxytocin genes in rats occurs in mutually exclusive sets of hypothalamic neurons. FEBS Lett 242, 144–148. 10.1016/0014-5793(88)81003-2.

25. Sullivan, G.M., Apergis, J., Bush, D.E.A., Johnson, L.R., Hou, M., and Ledoux, J.E. (2004). Lesions in the bed nucleus of the stria terminalis disrupt corticosterone and freezing responses elicited by a contextual but not by a specific cue-conditioned fear stimulus. Neuroscience 128, 7–14. 10.1016/j.neuroscience.2004.06.015.

26. Duvarci, S., Bauer, E.P., and Pare, D. (2009). The Bed Nucleus of the Stria Terminalis Mediates Inter-individual Variations in Anxiety and Fear. Journal of Neuroscience 29, 10357–10361. 10.1523/JNEUROSCI.2119-09.2009.

27. Moaddab, M., and Dabrowska, J. (2017). Oxytocin receptor neurotransmission in the dorsolateral bed nucleus of the stria terminalis facilitates the acquisition of cued fear in the fear-potentiated startle paradigm in rats. Neuropharmacology 121, 130–139. 10.1016/j.neuropharm.2017.04.039.

28. Goode, T.D., Ressler, R.L., Acca, G.M., Miles, O.W., and Maren, S. (2019). Bed nucleus of the stria terminalis regulates fear to unpredictable threat signals. Elife 8. 10.7554/eLife.46525.

29. DiBenedictis, B.T., Nussbaum, E.R., Cheung, H.K., and Veenema, A.H. (2017). Quantitative mapping reveals age and sex differences in vasopressin, but not oxytocin, immunoreactivity in the rat social behavior neural network. J Comp Neurol 525, 2549–2570. 10.1002/cne.24216.

30. Dabrowska, J., Hazra, R., Ahern, T.H., Guo, J.-D., McDonald, A.J., Mascagni, F., Muller, J.F., Young, L.J., and Rainnie, D.G. (2011). Neuroanatomical evidence for reciprocal regulation of the corticotrophin-releasing factor and oxytocin systems in the hypothalamus and the bed nucleus of the stria terminalis of the rat: Implications for balancing stress and affect. Psychoneuroendocrinology 36, 1312–1326. 10.1016/j.psyneuen.2011.03.003.

31. Duque-Wilckens, N., Steinman, M.Q., Laredo, S.A., Hao, R., Perkeybile, A.M., Bales, K.L., and Trainor, B.C. (2016). Inhibition of vasopressin V1a receptors in the medioventral bed nucleus of the stria terminalis has sex- and context-specific anxiogenic effects. Neuropharmacology 110, 59–68. 10.1016/j.neuropharm.2016.07.018.

32. Luo, P.X., Zakharenkov, H.C., Torres, L.Y., Rios, R.A., Gegenhuber, B., Black, A.M., Xu, C.K., Minie, V.A., Tran, A.M., Tollkuhn, J., et al. (2022). Oxytocin receptor behavioral effects and cell types in the bed nucleus of the stria terminalis. Horm Behav 143, 105203. 10.1016/j.yhbeh.2022.105203.

33. Dabrowska, J., Hazra, R., Ahern, T.H., Guo, J.-D., McDonald, A.J., Mascagni, F., Muller, J.F., Young, L.J., and Rainnie, D.G. (2011). Neuroanatomical evidence for reciprocal regulation of the corticotrophin-releasing factor and oxytocin systems in the hypothalamus and the bed nucleus of the stria terminalis of the rat: Implications for balancing stress and affect. Psychoneuroendocrinology 36, 1312–1326. 10.1016/j.psyneuen.2011.03.003.

34. Knobloch, H.S., Charlet, A., Hoffmann, L.C., Eliava, M., Khrulev, S., Cetin, A.H., Osten, P., Schwarz, M.K., Seeburg, P.H., Stoop, R., et al. (2012). Evoked axonal oxytocin release in the central amygdala attenuates fear response. Neuron 73, 553–566. 10.1016/j.neuron.2011.11.030.

35. Dabrowska, J., Hazra, R., Guo, J.-D., Li, C., DeWitt, S., Xu, J., Lombroso, P.J., and Rainnie, D.G. (2013). Striatal-Enriched Protein Tyrosine Phosphatase—STEPs Toward Understanding Chronic Stress-Induced Activation of Corticotrophin Releasing Factor Neurons in the Rat Bed Nucleus of the Stria Terminalis. Biological Psychiatry 74, 817–826. 10.1016/j.biopsych.2013.07.032.

36. Dabrowska, J., Hazra, R., Guo, J.-D., DeWitt, S., and Rainnie, D.G. (2013). Central CRF neurons are not created equal: phenotypic differences in CRF-containing neurons of the rat paraventricular hypothalamus and the bed nucleus of the stria terminalis. Front. Neurosci. 7. 10.3389/fnins.2013.00156.

37. Hammack, S.E., Mania, I., and Rainnie, D.G. (2007). Differential Expression of Intrinsic Membrane Currents in Defined Cell Types of the Anterolateral Bed Nucleus of the Stria Terminalis. Journal of Neurophysiology 98, 638–656. 10.1152/jn.00382.2007.

38. Rodríguez-Sierra, O.E., Turesson, H.K., and Pare, D. (2013). Contrasting distribution of physiological cell types in different regions of the bed nucleus of the stria terminalis. J. Neurophysiol. 110, 2037–2049. 10.1152/jn.00408.2013.

39. Martinon, D., Lis, P., Roman, A.N., Tornesi, P., Applebey, S.V., Buechner, G., Olivera, V., and Dabrowska, J. (2019). Oxytocin receptors in the dorsolateral bed nucleus of the stria terminalis (BNST) bias fear learning toward temporally predictable cued fear. Transl Psychiatry 9, 140. 10.1038/s41398-019-0474-x.

40. Francesconi, W., Berton, F., Olivera-Pasilio, V., and Dabrowska, J. (2021). Oxytocin excites BNST interneurons and inhibits BNST output neurons to the central amygdala. Neuropharmacology 192, 108601. 10.1016/j.neuropharm.2021.108601.

41. Althammer, F., Roy, R.K., Lefevre, A., Najjar, R.S., Schoenig, K., Bartsch, D., Eliava, M., Feresin, R.G., Hammock, E.A.D., Murphy, A.Z., et al. (2022). Altered PVN-to-CA2 hippocampal oxytocin pathway and reduced number of oxytocin-receptor expressing astrocytes in heart failure rats. J Neuroendocrinol 34, e13166. 10.1111/jne.13166.

42. Iwasaki, M., Lefevre, A., Althammer, F., Clauss Creusot, E., Łąpieś, O., Petitjean, H., Hilfiger, L., Kerspern, D., Melchior, M., Küppers, S., et al. (2023). An analgesic pathway from parvocellular oxytocin neurons to the periaqueductal gray in rats. Nat Commun 14, 1066. 10.1038/s41467-023-36641-7.

43. Pomrenze, M.B., Millan, E.Z., Hopf, F.W., Keiflin, R., Maiya, R., Blasio, A., Dadgar, J., Kharazia, V., De Guglielmo, G., Crawford, E., et al. (2015). A Transgenic Rat for Investigating the Anatomy and Function of Corticotrophin Releasing Factor Circuits. Front Neurosci 9, 487. 10.3389/fnins.2015.00487.

44. Zhang, L., Zetter, M.A., Hernández, V.S., Hernández-Pérez, O.R., Jáuregui-Huerta, F., Krabichler, Q., and Grinevich, V. (2024). Morphological Signatures of Neurogenesis and Neuronal Migration in Hypothalamic Vasopressinergic Magnocellular Nuclei of the Adult Rat. Int J Mol Sci 25, 6988. 10.3390/ijms25136988.

45. Millan, E.Z., Hopf, F.W., Keiflin, R., Maiya, R., Blasio, A., Dadgar, J., Kharazia, V., De Guglielmo, G., Crawford, E., Janak, P.H., et al. (2015). A Transgenic Rat for Investigating the Anatomy and Function of Corticotrophin Releasing Factor Circuits. Front. Neurosci. 9. 10.3389/fnins.2015.00487.

46. Pomrenze, M.B., Tovar-Diaz, J., Blasio, A., Maiya, R., Giovanetti, S.M., Lei, K., Morikawa, H., Hopf, F.W., and Messing, R.O. (2019). A Corticotropin Releasing Factor Network in the Extended Amygdala for Anxiety. J. Neurosci. 39, 1030–1043. 10.1523/JNEUROSCI.2143-18.2018.

47. Marir, R., Virsolvy, A., Wisniewski, K., Mion, J., Haddou, D., Galibert, E., Meraihi, Z., Desarménien, M.G., and Guillon, G. (2013). Pharmacological characterization of FE 201874, the first selective high affinity rat V _1A_ vasopressin receptor agonist: FE 201874, a selective rat V _1A_ agonist. Br J Pharmacol 170, 278–292. 10.1111/bph.12249.

48. Griffante, C., Green, A., Curcuruto, O., Haslam, C.P., Dickinson, B.A., and Arban, R. (2005). Selectivity of d[Cha4]AVP and SSR149415 at human vasopressin and oxytocin receptors: evidence that SSR149415 is a mixed vasopressin V1b/oxytocin receptor antagonist. Br J Pharmacol 146, 744–751. 10.1038/sj.bjp.0706383.

49. Manning, M., Misicka, A., Olma, A., Bankowski, K., Stoev, S., Chini, B., Durroux, T., Mouillac, B., Corbani, M., and Guillon, G. (2012). Oxytocin and Vasopressin Agonists and Antagonists as Research Tools and Potential Therapeutics: Oxytocin and vasopressin agonists and antagonists. Journal of Neuroendocrinology 24, 609–628. 10.1111/j.1365-2826.2012.02303.x.

50. Desai, N.S., Siegel, J.J., Taylor, W., Chitwood, R.A., and Johnston, D. (2015). MATLAB-based automated patch-clamp system for awake behaving mice. Journal of Neurophysiology 114, 1331–1345. 10.1152/jn.00025.2015.

51. Dabrowska, J., Martinon, D., Moaddab, M., and Rainnie, D.G. (2016). Targeting Corticotropin-Releasing Factor Projections from the Oval Nucleus of the Bed Nucleus of the Stria Terminalis Using Cell-Type Specific Neuronal Tracing Studies in Mouse and Rat Brain. J Neuroendocrinol 28. 10.1111/jne.12442.

52. Krashes, M.J., Koda, S., Ye, C., Rogan, S.C., Adams, A.C., Cusher, D.S., Maratos-Flier, E., Roth, B.L., and Lowell, B.B. (2011). Rapid, reversible activation of AgRP neurons drives feeding behavior in mice. J. Clin. Invest. 121, 1424–1428. 10.1172/JCI46229.

53. Knobloch, H.S., Charlet, A., Hoffmann, L.C., Eliava, M., Khrulev, S., Cetin, A.H., Osten, P., Schwarz, M.K., Seeburg, P.H., Stoop, R., et al. (2012). Evoked Axonal Oxytocin Release in the Central Amygdala Attenuates Fear Response. Neuron 73, 553–566. 10.1016/j.neuron.2011.11.030.

54. Paxinos, G., and Watson, C. (2009). The rat brain in stereotaxic coordinates Compact 6. ed. (Elsevier, Academic Press).

55. Yang, P., Wang, Z., Zhang, Z., Liu, D., Manolios, E.N., Chen, C., Yan, X., Zuo, W., and Chen, N. (2018). The extended application of The Rat Brain in Stereotaxic Coordinates in rats of various body weight. J Neurosci Methods 307, 60–69. 10.1016/j.jneumeth.2018.06.026.

56. Ben-Barak, Y., Russell, J.T., Whitnall, M.H., Ozato, K., and Gainer, H. (1985). Neurophysin in the hypothalamo-neurohypophysial system. I. Production and characterization of monoclonal antibodies. J Neurosci 5, 81–97. 10.1523/JNEUROSCI.05-01-00081.1985.

57. Williford, K.M., Taylor, A., Melchior, J.R., Yoon, H.J., Sale, E., Negasi, M.D., Adank, D.N., Brown, J.A., Bedenbaugh, M.N., Luchsinger, J.R., et al. (2023). BNST PKCδ neurons are activated by specific aversive conditions to promote anxiety-like behavior. Neuropsychopharmacol. 10.1038/s41386-023-01569-5.

58. Daniel, S.E., and Rainnie, D.G. (2016). Stress Modulation of Opposing Circuits in the Bed Nucleus of the Stria Terminalis. Neuropsychopharmacol 41, 103–125. 10.1038/npp.2015.178.

59. Schneider, C.A., Rasband, W.S., and Eliceiri, K.W. (2012). NIH Image to ImageJ: 25 years of image analysis. Nat Methods 9, 671–675. 10.1038/nmeth.2089.

60. Zaelzer, C., Gizowski, C., Salmon, C.K., Murai, K.K., and Bourque, C.W. (2018). Detection of activity-dependent vasopressin release from neuronal dendrites and axon terminals using sniffer cells. Journal of Neurophysiology 120, 1386–1396. 10.1152/jn.00467.2017.

61. Thirouin, Z.S., and Bourque, C.W. (2021). Mechanism and function of phasic firing in vasopressin-releasing magnocellular neurosecretory cells. J Neuroendocrinol 33. 10.1111/jne.13048.

62. Walker, D., and Davis, M. (2002). Quantifying fear potentiated startle using absolute versus proportional increase scoring methods: implications for the neurocircuitry of fear and anxiety. Psychopharmacology 164, 318–328. 10.1007/s00213-002-1213-0.

63. Viviani, D., Charlet, A., van den Burg, E., Robinet, C., Hurni, N., Abatis, M., Magara, F., and Stoop, R. (2011). Oxytocin selectively gates fear responses through distinct outputs from the central amygdala. Science 333, 104–107. 10.1126/science.1201043.

64. Owen, S.F., Tuncdemir, S.N., Bader, P.L., Tirko, N.N., Fishell, G., and Tsien, R.W. (2013). Oxytocin enhances hippocampal spike transmission by modulating fast-spiking interneurons. Nature 500, 458–462. 10.1038/nature12330.

65. Hu, B., Boyle, C.A., and Lei, S. (2021). Activation of Oxytocin Receptors Excites Subicular Neurons by Multiple Signaling and Ionic Mechanisms. Cereb Cortex 31, 2402–2415. 10.1093/cercor/bhaa363.

66. Migita, K., Hori, N., Manako, J., Saito, R., Takano, Y., and Kamiya, H. (1998). Effects of arginine-vasopressin on neuronal interaction from the area postrema to the nucleus tractus solitarii in rat brain slices. Neurosci Lett 256, 45–48. 10.1016/s0304-3940(98)00753-8.

67. Boyle, C.A., Hu, B., Quaintance, K.L., and Lei, S. (2021). Involvement of TRPC5 channels, inwardly rectifying K+ channels, PLCβ and PIP2 in vasopressin-mediated excitation of medial central amygdala neurons. J Physiol 599, 3101–3119. 10.1113/JP281260.

68. Viviani, D., Charlet, A., van den Burg, E., Robinet, C., Hurni, N., Abatis, M., Magara, F., and Stoop, R. (2011). Oxytocin Selectively Gates Fear Responses Through Distinct Outputs from the Central Amygdala. Science 333, 104–107. 10.1126/science.1201043.

69. Manning, M., Misicka, A., Olma, A., Bankowski, K., Stoev, S., Chini, B., Durroux, T., Mouillac, B., Corbani, M., and Guillon, G. (2012). Oxytocin and vasopressin agonists and antagonists as research tools and potential therapeutics. J. Neuroendocrinol. 24, 609–628. 10.1111/j.1365-2826.2012.02303.x.

70. Song, Z., Borland, J.M., Larkin, T.E., O’Malley, M., and Albers, H.E. (2016). Activation of oxytocin receptors, but not arginine-vasopressin V1a receptors, in the ventral tegmental area of male Syrian hamsters is essential for the reward-like properties of social interactions. Psychoneuroendocrinology 74, 164–172. 10.1016/j.psyneuen.2016.09.001.

71. Waltenspühl, Y., Ehrenmann, J., Vacca, S., Thom, C., Medalia, O., and Plückthun, A. (2022). Structural basis for the activation and ligand recognition of the human oxytocin receptor. Nat Commun 13, 4153. 10.1038/s41467-022-31325-0.

72. Gimpl, G., and Fahrenholz, F. (2001). The oxytocin receptor system: structure, function, and regulation. Physiol Rev 81, 629–683. 10.1152/physrev.2001.81.2.629.

73. Sala, M., Braida, D., Lentini, D., Busnelli, M., Bulgheroni, E., Capurro, V., Finardi, A., Donzelli, A., Pattini, L., Rubino, T., et al. (2011). Pharmacologic rescue of impaired cognitive flexibility, social deficits, increased aggression, and seizure susceptibility in oxytocin receptor null mice: a neurobehavioral model of autism. Biol Psychiatry 69, 875–882. 10.1016/j.biopsych.2010.12.022.

74. Neumann, I.D., and Landgraf, R. (2012). Balance of brain oxytocin and vasopressin: implications for anxiety, depression, and social behaviors. Trends Neurosci. 35, 649–659. 10.1016/j.tins.2012.08.004.

75. Chini, B., Mouillac, B., Ala, Y., Balestre, M.N., Trumpp-Kallmeyer, S., Hoflack, J., Elands, J., Hibert, M., Manning, M., and Jard, S. (1995). Tyr115 is the key residue for determining agonist selectivity in the V1a vasopressin receptor. EMBO J 14, 2176–2182. 10.1002/j.1460-2075.1995.tb07211.x.

76. Song, Z., and Albers, H.E. (2018). Cross-talk among oxytocin and arginine-vasopressin receptors: Relevance for basic and clinical studies of the brain and periphery. Front Neuroendocrinol 51, 14–24. 10.1016/j.yfrne.2017.10.004.

77. Lowbridge, J., Manning, M., Haldar, J., and Sawyer, W.H. (1977). Synthesis and some pharmacological properties of [4-threonine, 7-glycine]oxytocin, [1-(L-2-hydroxy-3-mercaptopropanoic acid), 4-threonine, 7-glycine]oxytocin (hydroxy[Thr4, Gly7]oxytocin), and [7-Glycine]oxytocin, peptides with high oxytocic-antidiuretic selectivity. J Med Chem 20, 120–123. 10.1021/jm00211a025.

78. Busnelli, M., Bulgheroni, E., Manning, M., Kleinau, G., and Chini, B. (2013). Selective and potent agonists and antagonists for investigating the role of mouse oxytocin receptors. J Pharmacol Exp Ther 346, 318–327. 10.1124/jpet.113.202994.

79. Gillard, E.R., Coburn, C.G., de Leon, A., Snissarenko, E.P., Bauce, L.G., Pittman, Q.J., Hou, B., and Currás-Collazo, M.C. (2007). Vasopressin autoreceptors and nitric oxide-dependent glutamate release are required for somatodendritic vasopressin release from rat magnocellular neuroendocrine cells responding to osmotic stimuli. Endocrinology 148, 479–489. 10.1210/en.2006-0995.

80. Goodson, J.L., and Thompson, R.R. (2010). Nonapeptide mechanisms of social cognition, behavior and species-specific social systems. Curr Opin Neurobiol 20, 784–794. 10.1016/j.conb.2010.08.020.

81. Lukas, M., and Neumann, I.D. (2013). Oxytocin and vasopressin in rodent behaviors related to social dysfunctions in autism spectrum disorders. Behav Brain Res 251, 85–94. 10.1016/j.bbr.2012.08.011.

82. Veinante, P., and Freund-Mercier, M.J. (1997). Distribution of oxytocin- and vasopressin-binding sites in the rat extended amygdala: a histoautoradiographic study. J. Comp. Neurol. 383, 305– 325.

83. Francesconi, W., Berton, F., Olivera-Pasilio, V., and Dabrowska, J. (2021). Oxytocin excites BNST interneurons and inhibits BNST output neurons to the central amygdala. Neuropharmacology 192, 108601. 10.1016/j.neuropharm.2021.108601.

84. Grundwald, N.J., Benítez, D.P., and Brunton, P.J. (2016). Sex-Dependent Effects of Prenatal Stress on Social Memory in Rats: A Role for Differential Expression of Central Vasopressin-1a Receptors. J. Neuroendocrinol. 28. 10.1111/jne.12343.

85. Nair, H.P., Gutman, A.R., Davis, M., and Young, L.J. (2005). Central oxytocin, vasopressin, and corticotropin-releasing factor receptor densities in the basal forebrain predict isolation potentiated startle in rats. J Neurosci 25, 11479–11488. 10.1523/JNEUROSCI.2524-05.2005.

86. Luo, P.X., Zakharenkov, H.C., Torres, L.Y., Rios, R.A., Gegenhuber, B., Black, A.M., Xu, C.K., Minie, V.A., Tran, A.M., Tollkuhn, J., et al. (2022). Oxytocin receptor behavioral effects and cell types in the bed nucleus of the stria terminalis. Hormones and Behavior 143, 105203. 10.1016/j.yhbeh.2022.105203.

87. Hasan, M.T., Althammer, F., Silva da Gouveia, M., Goyon, S., Eliava, M., Lefevre, A., Kerspern, D., Schimmer, J., Raftogianni, A., Wahis, J., et al. (2019). A Fear Memory Engram and Its Plasticity in the Hypothalamic Oxytocin System. Neuron 103, 133–146.e8. 10.1016/j.neuron.2019.04.029.

88. Rasie Abdullahi, P., Eskandarian, S., Ghanbari, A., and Rashidy-Pour, A. (2018). Oxytocin receptor antagonist atosiban impairs consolidation, but not reconsolidation of contextual fear memory in rats. Brain Research 1695, 31–36. 10.1016/j.brainres.2018.05.034.

89. Fam, J., Holmes, N., Delaney, A., Crane, J., and Westbrook, R.F. (2018). Oxytocin receptor activation in the basolateral complex of the amygdala enhances discrimination between discrete cues and promotes configural processing of cues. Psychoneuroendocrinology 96, 84–92. 10.1016/j.psyneuen.2018.06.006.

90. Missig, G., Ayers, L.W., Schulkin, J., and Rosen, J.B. (2010). Oxytocin Reduces Background Anxiety in a Fear-Potentiated Startle Paradigm. Neuropsychopharmacol 35, 2607–2616. 10.1038/npp.2010.155.

91. Ayers, L.W., Missig, G., Schulkin, J., and Rosen, J.B. (2011). Oxytocin Reduces Background Anxiety in a Fear-Potentiated Startle Paradigm: Peripheral vs Central Administration. Neuropsychopharmacol 36, 2488–2497. 10.1038/npp.2011.138.

92. Ayers, L., Agostini, A., Schulkin, J., and Rosen, J.B. (2016). Effects of oxytocin on background anxiety in rats with high or low baseline startle. Psychopharmacology 233, 2165–2172. 10.1007/s00213-016-4267-0.

93. Olivera-Pasilio, V., and Dabrowska, J. (2023). Fear-Conditioning to Unpredictable Threats Reveals Sex and Strain Differences in Rat Fear-Potentiated Startle (FPS). Neuroscience 530, 108–132. 10.1016/j.neuroscience.2023.08.030.

94. Neumann, I.D., and Slattery, D.A. (2016). Oxytocin in General Anxiety and Social Fear: A Translational Approach. Biol. Psychiatry 79, 213–221. 10.1016/j.biopsych.2015.06.004.

95. Walker, D.L., and Davis, M. (2008). Role of the extended amygdala in short-duration versus sustained fear: a tribute to Dr. Lennart Heimer. Brain Struct Funct 213, 29–42. 10.1007/s00429-008-0183-3.

96. Davis, M., Walker, D.L., Miles, L., and Grillon, C. (2010). Phasic vs Sustained Fear in Rats and Humans: Role of the Extended Amygdala in Fear vs Anxiety. Neuropsychopharmacol 35, 105– 135. 10.1038/npp.2009.109.

97. Yamauchi, N., Takahashi, D., Sugimura, Y.K., Kato, F., Amano, T., and Minami, M. (2018). Activation of the neural pathway from the dorsolateral bed nucleus of the stria terminalis to the central amygdala induces anxiety-like behaviors. Eur. J. Neurosci. 48, 3052–3061. 10.1111/ejn.14165.

98. Fodor, A., Kovács, K.B., Balázsfi, D., Klausz, B., Pintér, O., Demeter, K., Daviu, N., Rabasa, C., Rotllant, D., Nadal, R., et al. (2016). Depressive- and anxiety-like behaviors and stress-related neuronal activation in vasopressin-deficient female Brattleboro rats. Physiol Behav 158, 100–111. 10.1016/j.physbeh.2016.02.041.

99. Stoehr, J.D., Cheng, S.W., and North, W.G. (1993). Homozygous Brattleboro rats display attenuated conditioned freezing responses. Neurosci Lett 153, 103–106. 10.1016/0304-3940(93)90087-2.

100. Ambrogi Lorenzini, C., Bucherelli, C., Giachetti, A., and Tassoni, G. (1988). Aversive conditioning of homozygous and heterozygous D.I. Brattleboro rats in the light-dark box. Physiol Behav 42, 439–445. 10.1016/0031-9384(88)90173-4.

101. Pitkow, L.J., Sharer, C.A., Ren, X., Insel, T.R., Terwilliger, E.F., and Young, L.J. (2001). Facilitation of affiliation and pair-bond formation by vasopressin receptor gene transfer into the ventral forebrain of a monogamous vole. J Neurosci 21, 7392–7396. 10.1523/JNEUROSCI.21-18-07392.2001.

102. Bielsky, I.F., Hu, S.-B., Szegda, K.L., Westphal, H., and Young, L.J. (2004). Profound impairment in social recognition and reduction in anxiety-like behavior in vasopressin V1a receptor knockout mice. Neuropsychopharmacology 29, 483–493. 10.1038/sj.npp.1300360.

103. Egashira, N., Tanoue, A., Matsuda, T., Koushi, E., Harada, S., Takano, Y., Tsujimoto, G., Mishima, K., Iwasaki, K., and Fujiwara, M. (2007). Impaired social interaction and reduced anxiety-related behavior in vasopressin V1a receptor knockout mice. Behav Brain Res 178, 123– 127. 10.1016/j.bbr.2006.12.009.

104. Appenrodt, E., Schnabel, R., and Schwarzberg, H. (1998). Vasopressin administration modulates anxiety-related behavior in rats. Physiol Behav 64, 543–547. 10.1016/s0031-9384(98)00119-x.

105. Bredewold, R., and Veenema, A.H. (2018). Sex differences in the regulation of social and anxiety-related behaviors: insights from vasopressin and oxytocin brain systems. Curr. Opin. Neurobiol. 49, 132–140. 10.1016/j.conb.2018.02.011.

106. Whylings, J., Rigney, N., Peters, N.V., de Vries, G.J., and Petrulis, A. (2020). Sexually dimorphic role of BNST vasopressin cells in sickness and social behavior in male and female mice. Brain, Behavior, and Immunity 83, 68–77. 10.1016/j.bbi.2019.09.015.

107. Whylings, J., Rigney, N., de Vries, G.J., and Petrulis, A. (2021). Reduction in vasopressin cells in the suprachiasmatic nucleus in mice increases anxiety and alters fluid intake. Horm Behav 133, 104997. 10.1016/j.yhbeh.2021.104997.

108. Busnelli, M., and Chini, B. (2018). Molecular Basis of Oxytocin Receptor Signalling in the Brain: What We Know and What We Need to Know. Curr Top Behav Neurosci 35, 3–29. 10.1007/7854_2017_6.

109. Marir, R., Virsolvy, A., Wisniewski, K., Mion, J., Haddou, D., Galibert, E., Meraihi, Z., Desarménien, M.G., and Guillon, G. (2013). Pharmacological characterization of FE 201874, the first selective high affinity rat V1A vasopressin receptor agonist. Br J Pharmacol 170, 278–292. 10.1111/bph.12249.

110. Pena, A., Murat, B., Trueba, M., Ventura, M.A., Wo, N.C., Szeto, H.H., Cheng, L.L., Stoev, S., Guillon, G., and Manning, M. (2007). Design and synthesis of the first selective agonists for the rat vasopressin V(1b) receptor: based on modifications of deamino-[Cys1]arginine vasopressin at positions 4 and 8. J Med Chem 50, 835–847. 10.1021/jm060928n.

111. Manning, M., Misicka, A., Olma, A., Bankowski, K., Stoev, S., Chini, B., Durroux, T., Mouillac, B., Corbani, M., and Guillon, G. (2012). Oxytocin and vasopressin agonists and antagonists as research tools and potential therapeutics. J. Neuroendocrinol. 24, 609–628. 10.1111/j.1365-2826.2012.02303.x.

112. Serradeil-Le Gal, C., Wagnon, J., Garcia, C., Lacour, C., Guiraudou, P., Christophe, B., Villanova, G., Nisato, D., Maffrand, J.P., and Le Fur, G. (1993). Biochemical and pharmacological properties of SR 49059, a new, potent, nonpeptide antagonist of rat and human vasopressin V1a receptors. J Clin Invest 92, 224–231. 10.1172/JCI116554.

113. Griffante, C., Green, A., Curcuruto, O., Haslam, C.P., Dickinson, B.A., and Arban, R. (2005). Selectivity of d[Cha4]AVP and SSR149415 at human vasopressin and oxytocin receptors: evidence that SSR149415 is a mixed vasopressin V1b/oxytocin receptor antagonist. Br J Pharmacol 146, 744–751. 10.1038/sj.bjp.0706383.

114. Allen Institute for Brain Science. (2024). Allen Mouse Brain Atlas (2004). Retrieved from https://mouse.brain-map.org (Accessed March 30, 2024). http://atlas.brain-map.org/atlas?atlas=1#atlas=1&plate=100960312&structure=554&x=5222.879464285715&y=3571.841212681362&zoom=-3&resolution=11.97&z=6

